# RNA Helix Thermodynamics: The End Game

**DOI:** 10.1101/2021.10.16.464667

**Authors:** Jeffrey Zuber, Susan J. Schroeder, Hongying Sun, Douglas H. Turner, David H. Mathews

## Abstract

Nearest neighbor parameters for estimating the folding stability of RNA secondary structures are in widespread use. For helices, current parameters penalize terminal AU base pairs relative to terminal GC base pairs. We curated an expanded database of helix stabilities determined by optical melting experiments. Analysis of the updated database shows that terminal penalties depend on the sequence identity of the adjacent penultimate base pair. New nearest neighbor parameters that include this additional sequence dependence accurately predict the measured values of 271 helices in an updated database with a correlation coefficient of 0.982. This refined understanding of helix ends facilitates fitting terms for base pair stacks with GU pairs. Prior parameter sets treated 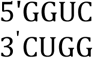 separately from other 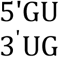 stacks. The improved understanding of helix end stability, however, makes the separate treatment unnecessary. Introduction of the additional terms was tested with three optical melting experiments. The average absolute difference between measured and predicted free energy changes at 37° C for these three duplexes containing terminal adjacent AU and GU pairs improved from 1.38 to 0.27 kcal/mol. This confirms the need for the additional sequence dependence in the model.

## INTRODUCTION

Over 80% of the human genome is transcribed into RNA, but less than 3% of the RNA codes for proteins (1,2). Functions for most RNA in the biosphere are still being discovered but already include catalysis (3), control of transcription, translation and expression (4-6), templating for synthesis of DNA (7) and RNA (8), recognition of sites for modification and editing (9-11) and sometimes combining such functions (12). RNA is the genomic material for many viruses, including human pathogens such as SARS and SARS-CoV-2, influenza, HIV, Ebola, and Hepatitis C. RNA can also be the basis for vaccines against some of these viruses. For example, mRNA vaccines show 94-95% effectiveness against SARS-CoV-2 infections (13).

RNA sequence determines the base pairing and 3D structure as well as function of the RNA. Prediction of secondary structure, i.e. the canonical set of Watson-Crick-Franklin (WCF) and GU base pairs, from sequence is a first step in predicting 3D structure (14) and in finding RNAs with common structures and functions (15,16). Some RNA, such as riboswitches, have more than one structure, and the ability to change structure is critical to function (5).

Secondary structure can be predicted from one or more sequences by minimizing free energy change for folding, ΔG°, often augmented with information from chemical mapping and/or sequence comparison. Usually, about half the nucleotides in an RNA are canonically paired (17). GU pairs play important roles in RNA structure and function as sites for binding metal ions (18,19), therapeutics (20), proteins, or metabolites (21).

A database of thermodynamic measurements for helices with canonical pairs and model non-canonical motifs forms the foundation for folding free energy predictions of RNA structure. These data are then fit to a nearest neighbor (NN) model to estimate parameters that can be used to predict folding stabilities of any RNA secondary structure (22). Hallmarks of the NN model are that each stability increment depends on local sequence and that total stability is the sum of the increments.

The model and parameters for approximating stabilities of WCF base-paired helixes have not changed substantially since 1998 (23,24). Individual parameters for nearest neighbors containing at least one GU pair, however, were revised on the basis of new data (24). In that revision, the penalty of 0.45 kcal/mol applied to terminal AU pairs and previously assumed for terminal GU pairs (23), was found unnecessary for GU pairs. Expansion of the database for duplexes, particularly those with terminal GU pairs (25) and the data presented here, make possible more extensive considerations of terminal effects on base pair stability. In particular, the data allow expansion of the model to include six new parameters specific for terminal nearest neighbors. Surprisingly, stabilities of terminal GU and AU pairs depend on whether the neighboring pair is a GC, AU, or GU pair. Incorporating this effect in the NN model also produces significant revision of parameters for internal 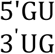 and 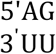 nearest neighbors. With these Changes, 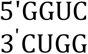 fits the NN model rather than being an outlier as considered previously (26). Thus, the new model presented here will be especially important for predicting structures containing GU pairs.

It is not surprising that GU pairs are more idiosyncratic than WCF pairs. Guanine has more hydrogen bonding groups and a larger dipole moment than other bases (27). Base stacking and hydrogen bonding that stabilize GU pairs can vary depending on local context, including position in a helix. Base stacking depends on interactions with both bases of a nearest neighbor. GU pairs can adopt different hydrogen bonded configurations and stacking interactions (Figure 1). In Figure 1A, the terminal UG pair is in a 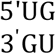 nearest neighbor and has a single hydrogen bond while the penultimate GU pair has two hydrogen bonds. Conformation of the terminal UG pair may be influenced by solvent interactions or crystal contacts through stacking interactions with the terminal UG pair of an adjacent molecule. In Figure 1B, the 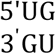 nearest neighbor is flanked by WCF GC pairs on both sides in the middle of the helix, i.e. 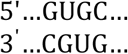. In that context, both the UG and GU pair have a hydrogen bond from the G carbonyl to the U imino proton, a bifurcated hydrogen bond between the U carbonyl at C2 and the G imino and amino protons, and extensive cross-strand guanine stacking. In contrast, when the sequence is reversed, i.e. 5’…CGUG… compared to 5’…GUGC…, in the self-complementary duplex (Figure 1C), each GU pair has a single bifurcated hydrogen bond, and there is no cross-strand stacking.

**Figure 1:**
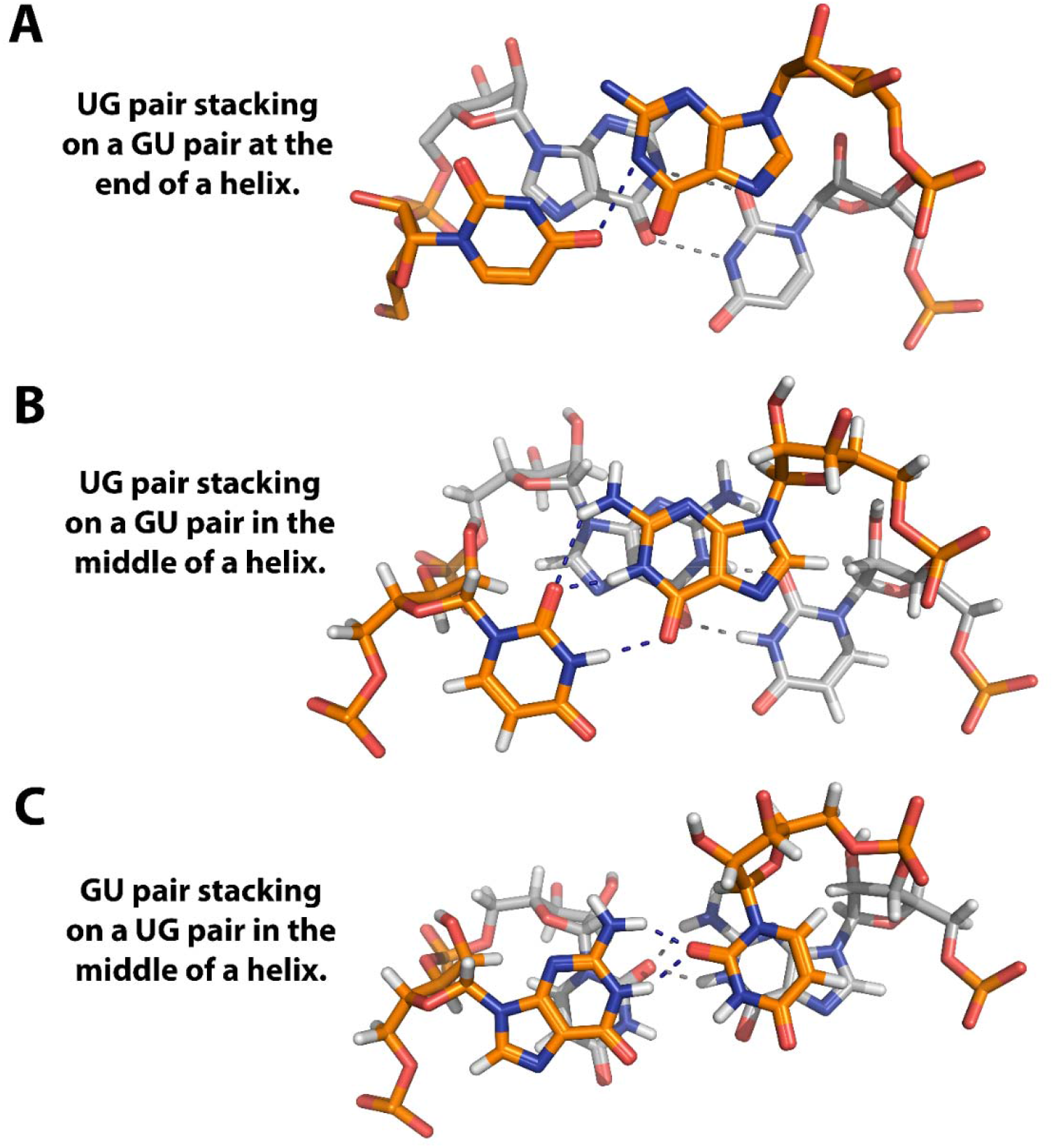
GU pair stacking and hydrogen bonding for three contexts of tandem GU pairs. A. X-ray crystal structure (1.4 Å resolution, R_free_ = 20.7%) with terminal stacking GU pairs in 5’**UG**CUCCUAGUACGUAAGGACCGGAG**UG**, PDB ID# 1MSY (124). Nucleotides in bold in the sequence are shown. The nucleotides with gold carbon atoms are in the forefront and these constitute the terminal base pair, while nucleotides with gray carbon atoms are in the back. The crystal packing has the terminal GU pairs of two molecules stacking on each other. B. NMR structure with internal UG pairs in (5’GAG**UG**CUC)_2_, PDB ID# 1EKA (116). 28 unique NOE measurements define the orientation for these nucleotides. C. NMR structure with internal GU pairs in (5’GGC **GU**GCC)_2_. PDB ID# 1EKD (116). 26 unique NOE measurements define the orientation for these nucleotides.

Reported thermodynamic stabilities for the internal NN GU stacks in Figures 1B and C reflect the different nucleotide configurations and effect of considering terminal effects. The 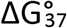 of 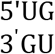 and 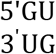 motifs are -0.38 and -0.19 kcal/mol, respectively, in the new model as compared to -0.57 and +0.72 kcal/mol in a prior model (24). The new parameters also include an increment of -0.74 kcal/mol for terminal consecutive GU pairs, i.e. a GU end on a GU pair, like those shown in Figure 1A. In contrast, previous models did not add favorable folding free energy for this sequence motif. In many NMR structures with terminal GU and AU pairs, these pairs show more dynamic behavior relative to internal pairs. This is observed as broad imino proton resonances and fewer NOE restraints (25,28-32). Thus, sequence orientation, stacking interactions, hydrogen bonding, and nucleotide dynamics are important factors in the structure and stabilities of GU pairs. They are now more accurately accounted for in the new thermodynamic parameter set. This should improve predictions of secondary structure from sequence.

## MATERIALS AND METHODS

### Optical Melting Experiment Database

For this analysis, optical melting experiments were compiled through an extensive literature review (24,25,28,33-61). Enumeration of all melting experiments included in this analysis is available in Supplemental Table 1 and in the spreadsheet provided in the supplementary materials. Experiments are included if only unmodified nucleotides are present and buffer has 1 M Na+ with pH between 6.5 and 7.5. Additionally, values of ΔH° determined with van’t Hoff plots and curve fitting of absorption vs temperature had to agree within 15% (42), which is consistent with an approximate 2-state transition.

Most melting experiments for generating NN parameters have used 1 M Na^+^. This was initially chosen because available duplexes had many AU pairs and therefore needed high salt to melt in convenient temperature ranges (62,63). The most important result from thermodynamic studies is the relative sequence dependence of nearest neighbor stability. This is expected not to depend on salt conditions because there is no site binding of Na^+^ to fully base paired RNA. Local concentrations of mobile cations around large folded RNAs, however, depend on the local charge density of phosphate groups. Manning developed a first order cation condensation model that predicts local concentrations of cations around RNA do not depend on bulk concentrations (64). For A-form double helical and single strand RNA, respectively, the local “ion atmosphere” is predicted to have 1.7 M and 0.4 M of M^+^ ions (65). They respectively neutralize 0.8 and 0.6 of the phosphate charge. Mg^2+^ will also bind as part of the ion atmosphere and neutralize 0.9 and 0.8, respectively of backbone charge. More detailed computations and experiments agree qualitatively with expectations from Manning theory (28,51,57,66-71).

### Feature Correlations

Feature correlations were calculated for each model using the R statistical programming language (72). The resulting correlation matrices were then visualized with the *R corrplot* library (73).

### Fitting Linear Models

Parameter models were fit using measured 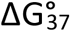 and ΔH° values for each optical melting experiment. For the fit of WCF stacking parameters, the theoretical contribution of RT ln(2) due to 2-fold symmetry of self-complementary duplexes, was subtracted from the experimentally measured duplex 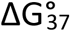 (23,74). For the fit of GU stacking parameters, the contributions due to sequence symmetry and the WCF stacks from each duplex with any GU base pairs were subtracted from measurements. The calculated 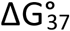 and ΔH° are then used to fit linear models in the R statistical programming language using the base function *lm (72)*. ΔS° values for the nearest neighbor parameters are calculated from the 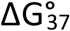 and ΔH° values.

To estimate uncertainty in NN parameter values, a covariation analysis was used to account for the dependencies (due to the nested nature of the regressions) and correlation (due to a base pair appearing in up to two neighboring stacks) between parameters (75) (76). To perform covariation analysis, the optical melting data were resampled within experimental error (ΔH_σ_= 12% ΔH° and 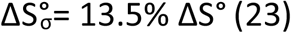) The resampling was performed with the *mvrnorm* function from the R MASS library (77), which preserves the observed correlation between ΔH° and ΔS° (ρ= 0.9996 (23)). The updated experimental values are then used to recalculate multiple sets of model parameters. The sets of model parameters are then used to calculate average values for each parameter as well as covariation (75,76). The standard errors of regression, which neglect the correlations and the effect of nested regressions, for the NN parameters can be found in Supplemental Table 2.

### Leave-One-Out Analysis

To assess the impact of any one experimental value on the fit models, models were fit in which each experimental value was individually excluded from the fitting data. The root mean square deviations (RMSDs) in parameter values were calculated from the model fit to the full data set to measure the impact of excluding each individual experimental value.

### Optical Melting Experiments

Optical melting experiments were conducted following standard protocols described in (78). Oligonucleotides were purchased from Integrated DNA Technologies including purification with standard desalting procedures and assessment of purity by mass spectrometry. Oligonucleotides were dissolved in milliQ water, and the absorbance at 260 nm at 80 °C was measured. The appropriate amount of oligonucleotide was dried in a speed vac and resuspended in standard melting buffer of 1 M NaCl, 20 mM sodium cacodylate, pH 7, and 0.5 mM Na_2_EDTA. Optical melting experiments were conducted in a Beckman DU800 UV-Vis spectrometer with a custom sample holder and cuvettes at 0.1 cm and 1.0 cm path lengths. Absorbance vs. temperature was measured at 260 and 280 nm. Data was analyzed at 280 nm with Meltwin software (51).

### Stacking Term Counts

An archive of RNA sequences of known secondary structure {Sloma, 2016 #1348} was analyzed to count the number of occurrences of each stacking parameter. A Python script was used to parse each structure into individual helices and then to parse each helix into component stacking and helix end parameters.

## Results

### AU End Parameters Depend on Penultimate Pair

Prior work demonstrated that multiple GU terminal base pairs impact the stability of helical duplexes (25), and this motivated a reexamination of the treatment of helix ends. New terms to account for the end of a helix were introduced into the NN model. This was done by including a parameter for an AU terminal pair on an AU penultimate pair (not accounting for orientation of the two pairs and therefore applying to 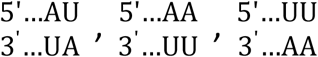, or 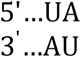 helix ends) and a parameter for an AU terminal pair on a CG penultimate pair (applying to 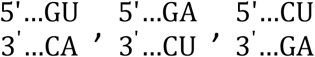, or 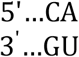 helix ends). These parameters are applied in addition to the base pair stacking parameter for these end stacks. This model is consistent with the observation that the thermodynamic stability bonus conferred by multiple GU terminal pairs is largely independent of orientation of the GU pairs (25).

Comparisons between the updated model and those used in the 1998 and 2004 NN models are in Table 1. Fitting to the model with the modified helix end parameters resulted in only moderate changes to the WCF stacks. The parameters for intermolecular initiation and the individual NN stacks were all within error of the 1998 parameters for both 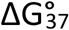and ΔH°. The only significant change was that an AU terminal pair on an AU penultimate pair is more favorable compared to the 1998 and 2004 models.

**Table 1A.** The 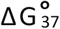 and for ΔH° nearest neighbor parameters for helices composed of WCF pairs.

| Feature | New Model<sup>§</sup> |  |  | 1998 Model<sup>¶</sup> |  |
| --- | --- | --- | --- | --- | --- |
| | $\Delta G_{37}^\circ$<br>(kcal/mol) | $\Delta H^\circ$<br>(kcal/mol) | $\Delta S^\circ$<br>(eu) | $\Delta G_{37}^\circ$<br>(kcal/mol) | $\Delta H^\circ$<br>(kcal/mol) |
| GC/CG | -3.46 ± 0.08 | -16.52 ± 1.57 | -42.13 ± 4.26 | -3.42 ± 0.08 | -14.88 ± 1.58 |
| CC/GG | -3.28 ± 0.08 | -13.94 ± 1.18 | -34.41 ± 3.58 | -3.26 ± 0.07 | -13.39 ± 1.24 |
| GA/CU | -2.42 ± 0.05 | -13.75 ± 1.00 | -36.53 ± 3.16 | -2.35 ± 0.06 | -12.44 ± 1.20 |
| CG/GC | -2.33 ± 0.09 | -9.61 ± 1.57 | -23.46 ± 4.74 | -2.36 ± 0.09 | -10.64 ± 1.65 |
| AC/UG | -2.25 ± 0.06 | -11.98 ± 1.17 | -31.37 ± 3.86 | -2.24 ± 0.06 | -11.40 ± 1.23 |
| CA/GU | -2.07 ± 0.07 | -10.47 ± 1.25 | -27.08 ± 3.73 | -2.11 ± 0.07 | -10.44 ± 1.28 |
| AG/UC | -2.01 ± 0.07 | -9.34 ± 1.23 | -23.66 ± 3.63 | -2.08 ± 0.06 | -10.48 ± 1.24 |
| UA/AU | -1.29 ± 0.08 | -9.16 ± 1.71 | -25.40 ± 5.55 | -1.33 ± 0.09 | -7.69 ± 2.02 |
| AU/UA | -1.09 ± 0.07 | -8.91 ± 1.55 | -25.22 ± 4.75 | -1.10 ± 0.08 | -9.38 ± 1.68 |
| AA/UU | -0.94 ± 0.04 | -7.44 ± 0.80 | -20.98 ± 2.56 | -0.93 ± 0.03 | -6.82 ± 0.79 |
| Initiation | +4.10 ± 0.24 | +4.66 ± 3.85 | 1.78 ± 11.93 | +4.09 ± 0.22 | +3.61 ± 4.12 |
| Symmetry | +0.43 | 0 | -1.38 | +0.43 | 0 |
| AU End on AU | +0.22 ± 0.06 | +4.36 ± 1.23 | 13.35 ± 3.83 | +0.45 ± 0.04<sup>‡</sup> | +3.72 ± 0.83<sup>‡</sup> |
| AU End on CG | +0.44 ± 0.04 | +3.17 ± 0.80 | 8.79 ± 2.50 | +0.45 ± 0.04<sup>‡</sup> | +3.72 ± 0.83<sup>‡</sup> |

**Table 1B.** The 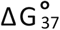 and for ΔH° nearest neighbor parameters for stacks with GU pairs.

| Feature | New Model<sup>§</sup> |  |  | 2012 Model<sup>*</sup> |  |
| --- | --- | --- | --- | --- | --- |
| | $\Delta G_{37}^\circ$ (37 °C)<br>(kcal/mol) | $\Delta H^\circ$<br>(kcal/mol) | $\Delta S^\circ$<br>(eu) | $\Delta G_{37}^\circ$ (37 °C)<br>(kcal/mol) | $\Delta H^\circ$<br>(kcal/mol) |
| GC/UG | -2.23 ± 0.07 | -14.73 ± 1.44 | -40.32 ± 4.60 | -2.15 ± 0.10 | -11.09 ± 1.78 |
| CU/GG | -1.93 ± 0.08 | -9.26 ± 1.58 | -23.64 ± 5.16 | -1.77 ± 0.09 | -9.44 ± 1.76 |
| GG/CU | -1.80 ± 0.07 | -12.41 ± 1.52 | -34.23 ± 4.72 | -1.80 ± 0.09 | -7.03 ± 1.75 |
| CG/GU | -1.05 ± 0.07 | -5.64 ± 1.47 | -14.83 ± 4.57 | -1.25 ± 0.09 | -5.56 ± 1.68 |
| AU/UG | -0.76 ± 0.07 | -9.23 ± 1.61 | -27.32 ± 5.09 | -0.90 ± 0.08 | -7.39 ± 1.65 |
| GA/UU | -0.60 ± 0.06 | -10.58 ± 1.52 | -32.19 ± 4.81 | -0.51 ± 0.08 | -10.38 ± 1.79 |
| UG/GU | -0.38 ± 0.07 | -8.76 ± 1.74 | -27.04 ± 5.21 | -0.57 ± 0.19 | -12.64 ± 4.01 |
| UA/GU | -0.22 ± 0.07 | -2.72 ± 1.54 | -8.08 ± 4.79 | -0.39 ± 0.09 | -0.96 ± 1.80 |
| GG/UU | -0.20 ± 0.08 | -9.06 ± 1.89 | -28.57 ± 6.04 | -0.25 ± 0.16 | -17.82 ± 3.75 |
| GU/UG | -0.19 ± 0.08 | -7.66 ± 1.80 | -24.11 ± 5.81 | +0.72 ± 0.19 | -13.83 ± 4.21 |
| AG/UU | +0.02 ± 0.06 | -5.10 ± 1.45 | -16.53 ± 4.56 | -0.35 ± 0.08 | -3.96 ± 1.73 |
| GGUC/CUGG | (-3.80 ± 0.13)<sup>†</sup> | (-32.49 ± 2.75)<sup>†</sup> | (-92.57 ± 11.09)<sup>†</sup> | -4.12 ± 0.54 | -30.80 ± 8.87 |
| AU End on GU | -0.71 ± 0.15 | 5.16 ± 2.99 | 18.96 ± 9.15 | +0.45 ± 0.04<sup>¶</sup> | +3.72 ± 0.83<sup>¶</sup> |
| GU End on CG | 0.13 ± 0.08 | 3.91 ± 1.43 | 12.17 ± 4.34 | 0.00 ± 0.00<sup>◊</sup> | 0.00 ± 0.00<sup>◊</sup> |
| GU End on AU | -0.31 ± 0.06 | 3.65 ± 1.37 | 12.78 ± 4.23 | 0.00 ± 0.00<sup>◊</sup> | 0.00 ± 0.00<sup>◊</sup> |
| GU End on GU | -0.74 ± 0.08 | 6.23 ± 2.12 | 22.47 ± 6.65 | 0.00 ± 0.00<sup>◊</sup> | 0.00 ± 0.00<sup>◊</sup> |

<sup>§</sup>Uncertainty values were calculated from the experiment covariation analysis.
Uncertainty values from standard errors of regression are listed in Supplemental Table 2.
<sup>¶</sup>Parameters taken from (23). <sup>‡</sup>The 1998 model did not have separate values for each AU
End variant. <sup>\*</sup>Values taken from (24). <sup>†</sup>The new model does not have this parameter. The
shown value is the result of combining the NN stacks for that sequence. <sup>o</sup>The 2012 model did not include a parameter for terminal GU base pairs.

The updated model shows excellent correlations between predicted and measured values of 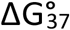 (R^2^=0.9823, Figure 2) and ΔH° (R^2^=0.8877, Supplemental Figure 1). The correlations between the model feature frequencies are modest and are mostly limited to expected correlations between stacks that can extend on each other (Supplemental Figure 2). The predicted folding 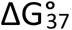 were within 0.5 kcal/mol of the measured value for 86.4% of the experiments (Supplemental Figure 3). Predicted ΔH° are within 5 kcal/mol of the measured value for 76% of the experiments (Supplemental Figure 3).

**Figure 2:**
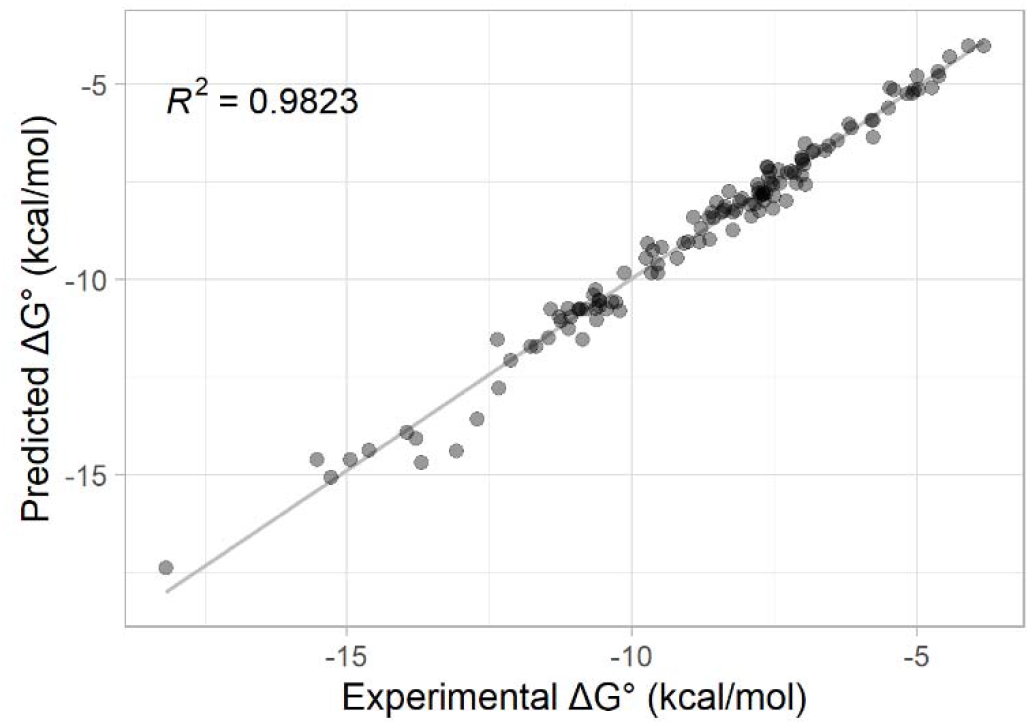
Correlation between predicted and measured 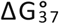 for duplexes with only WCF pairs. 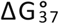 values predicted from updated nearest neighbor parameters for duplexes composed solely canonical WCFW-C-F base pairs in Table 1A plotted against values determined from optical melting experiments.

The impact of each optical melting experiment was determined by fitting the NN parameters on a data set that excluded that individual experiment and comparing the resulting parameter values to those fit on the full data set. The root mean squared deviations (RMSDs) in 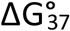 and ΔH° parameter values for these leave-one-out (LOO) data sets can be seen in Supplemental Figure 4. No one individual experiment heavily impacted the parameter values. The biggest impacts were RMSDs of 0.0363 kcal/mol in 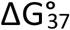 and 0.3346 kcal/mol in ΔH°, substantially smaller than uncertainty in the parameter values.

### GU Stacking Parameters

A similar model for terminal AU and GU stacks was used when fitting duplexes with GU base pairs. The model requires terms for an AU end with a penultimate GU pair, a terminal GU pair with a penultimate AU pair, a terminal GU pair with a penultimate GC pair, and a GU pair with a penultimate GU pair. The orientation of the two pairs is not considered.

Prior GU stack NN parameter sets treated 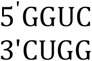 as a special, non-nearest neighbor case. When results from a fitting model including a parameter for the non-nearest neighbor quadruplet 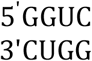, however, were compared to results for a model not including that parameter, the other parameter values were all within uncertainty of each other. Additionally, for each duplex containing 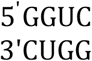 predicted 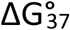 and ΔH° values from each model were also close to each other and to predicted 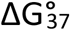 and ΔH° values from the 2012 model (24) (Supplemental Table 3). Additionally, the addition of the special 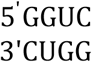 parameter results in almost identical R^2^ value for the fit (0.9256 vs 0.9267) (Figure 3 and Supplemental Figure 12). Evidently, a special, non-nearest neighbor parameter is not needed in the updated model when end effects for terminal AU and GU nearest neighbors are accounted for.

**Figure 3:**
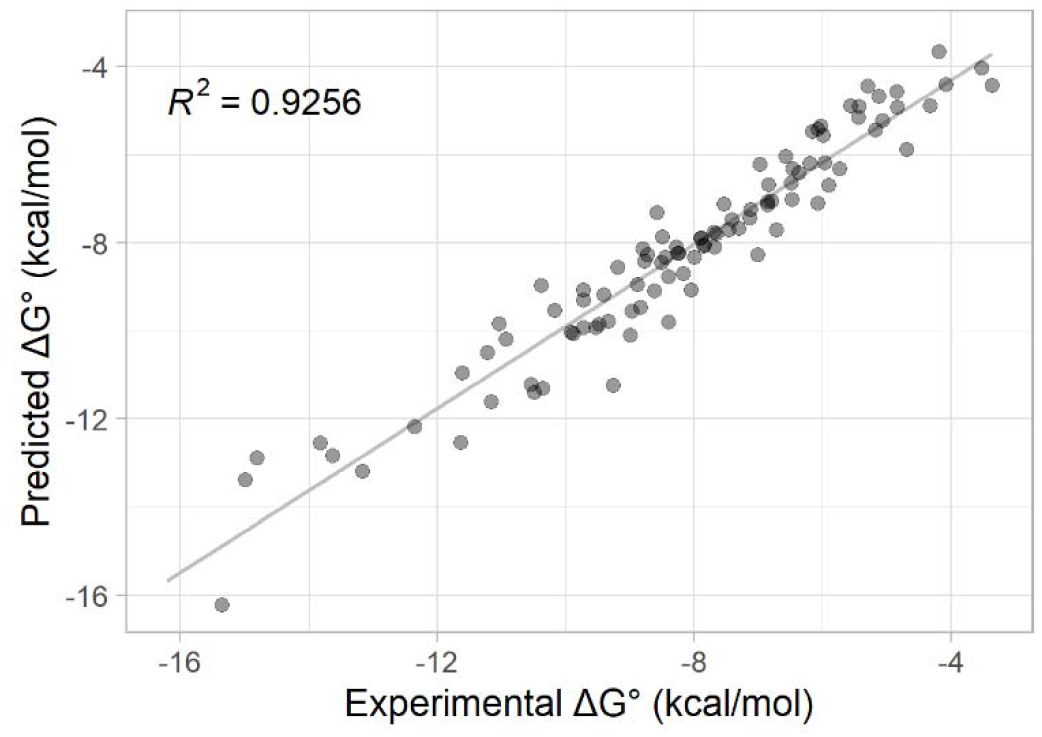
Correlation between predicted and observed 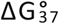 for duplexes with WCF and GU pairs. 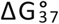 values predicted from parameters in Table 1 plotted against values determined from optical melting experiments.

For the GU internal stacking NN parameters, the most substantial change is for the 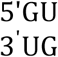 stack, where the 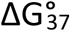 changed from +0.72 kcal/mol to -0.19 kcal/mol between the previous (24) and new models. An additional increment of -0.74 is added for a terminal 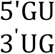 stack. The second most substantial change is for the 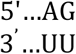 stack, which went from having a 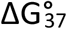 contribution of -0.35 kcal/mol (24) to +0.02 kcal/mol, but with an additional end increment of -0.31 kcal/mol.

The updated model shows good correlation between predicted and measured values for folding 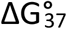 (R^2^=0.9256, Figure 3) and ΔH° (R^2^=0.7659, Supplemental Figure 5). Correlations between model feature frequencies are mostly limited to expected correlations between the GU on GU end feature and the three possible stacks that can form that end (Supplemental Figure 6). For folding 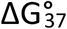, 53.1% of experiments had predicted values within 0.5 kcal/mol of the measured value (85.7% were within 1 kcal/mol) (Supplemental Figure 7). For ΔH°, 57.1% of experiments had predictions within 5 kcal/mol of the measured value (79.6% were within 10 kcal/mol) (Supplemental Figure 7).

RMSDs in 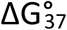 and ΔH° parameter values for the LOO data sets can be seen in Supplemental Figure 8. As with the WCF stacking parameters, no one individual experiment heavily impacted the parameter values. The biggest impacts are RMSDs of 0.0641 kcal/mol in 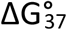 and 0.8675 kcal/mol in ΔH°, smaller than the uncertainty in the parameter values.

Uncertainty in parameter values for the updated NN model presented in Tables 1A and 1B were determined from a covariation analysis, which randomly perturbed the experimental values within experimental uncertainty and calculated the covariance matrix from observed changes in parameter values. Covariances between parameters are presented in Supplemental Figures 9 and 10. Covariances were generally small, with the strongest interactions between the intermolecular initiation parameter and parameters for individual stacks. There are also weaker interactions between GU end terms and equivalent internal GU stacks that can form that end term. For example, the GU on GU end parameter value is negatively correlated with the values for internal 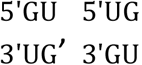 and 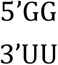 stacks.

### Additional Melting Experiments Support the Model

Three duplexes, designed to test features of the new model for thermodynamic parameters with terminal AU pairs and penultimate GU pairs, were studied by optical melting. A significant difference between the two models occurs for the end parameter for an AU end on a penultimate GU pair. The new and previous (23) models use values of -0.72 kcal/mol and +0.45 kcal/mol at 37 °C, respectively. All three duplexes in Table 2 contain this motif, which was represented in only two duplex sequences in the database of optical melting experiments. The duplexes, (5’UGUCGAUA)_2_ and (5’AUAGCUGU)_2_ differ in orientation of the terminal AU pair stacking on the penultimate GU pair. Duplex (5’AUUCGAGU)_2_ contains the motif 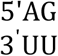, which has values of -0.03 and -0.35 kcal/mol for the new and prior models, respectively. Table 2 compares predictions based on the new and prior model with the experimentally measured thermodynamic values for these three duplexes. The experimental results confirm improvement of the new model.

**Table 2:** Melting buffer was 1 M NaCl, 20 mM sodium cacodylate pH 7, and 0.5 mM Na_2_EDTA. *a)* Predicted 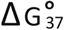 are calculated using Table 1 values for the new model parameters. *b)* Predicted 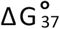 are calculated from the previous models described in (24) and (23). The duplex values in italics show borderline apparent 2-state behavior with a difference in ΔH° between the two analyses of 18.2%.

| Duplex | New Model<sup>a</sup> | Prior Model<sup>b</sup> | Van't Hoff Plot Analysis |  |  |  | Curve Fit Analysis |  |  |  |
| --- | --- | --- | --- | --- | --- | --- | --- | --- | --- | --- |
| | $-\Delta G_{37}^{\circ}$ (kcal/mol) | $-\Delta G_{37}^{\circ}$ (kcal/mol) | $-\Delta G_{37}^{\circ}$ (kcal/mol) | $-\Delta H^{\circ}$ (kcal/mol) | $-\Delta S^{\circ}$ (eu) | T<sub>m</sub> (°C)<sup>†</sup> | $-\Delta G_{37}^{\circ}$ (kcal/mol) | $-\Delta H^{\circ}$ (kcal/mol) | $-\Delta S^{\circ}$ (eu) | T<sub>m</sub> (°C)<sup>†</sup> |
| (5'UGUCGAUA)<sub>2</sub> | 6.02 ± 0.31 | 4.22 ± 0.37 | 6.10 ± 0.01 | 59.32 ± 0.01 | 171.58 ± 0.01 | 34.8 | 6.11 ± 0.18 | 66.83 ± 3.42 | 195.78 ± 11.21 | 39.0 |
| (5'AUAGCUGU)<sub>2</sub> | 6.33 ± 0.29 | 4.74 ± 0.37 | 5.81 ± 0.02 | 44.15 ± 0.01 | 123.63 ± 0.02 | 32.0 | 5.86 ± 0.26 | 47.88 ± 6.08 | 135.48 ± 19.18 | 38.2 |
| *(5'AUUCGAGU)<sub>2</sub>* | 5.54 ± 0.26 | 4.14 ± 0.36 | 5.32 ± 0.03 | 34.58 ± 0.01 | 94.33 ± 0.02 | 26.5 | *5.10 ± 0.29* | *42.30 ± 2.98* | *119.94 ± 9.69* | 32.9 |

<sup>†</sup> T<sub>m</sub> values were calculated for a concentration of 1×10<sup>-4</sup> M

## Discussion

The database of thermodynamic parameters forms the foundation for predictions of RNA structure and function in many widely used software suites (14,80-83,84,85-87). These RNA structure prediction programs enable design of mRNA vaccine sequences (13,88,89), analysis of metaproperties of transcriptomic changes in response to stress (90-92), determination of effects of nucleotide modifications on folding stability (93-95), discovery of accessible regions to target with antisense DNA or siRNA (96-99), and rational design of small molecules targeting RNA (100-103). Curation and improvement of the RNA thermodynamic database facilitates hypothesis-driven RNA research in many fields and has significant impact on the RNA community. The progress reported here expands, compiles, and presents the thermodynamic NN parameters for WCF and GU pairs. Statistical significance of the new parameters is robust. Inclusion of helix-end effects for AU and GU pairs improves predictions of helices with these common motifs and resolves previously poorly understood terms for “special cases” of motifs containing GU pairs.

This work presents the next advance in development of a robust NN model for predicting RNA duplex stabilities. The NN model for ΔG° of RNA helixes composed of canonical pairs uses stacks of adjacent base pairs (104-106). This assumes that the total ΔG° and temperature dependence, ΔH°, for helix formation can be approximated by summing ΔG°s and ΔH°s assigned to nearest neighbors of canonical pairs. The experimental foundation for this approach was laid by Uhlenbeck and Martin in the Doty lab when they used optical melting to measure thermodynamics of duplex formation (62,63). As a postdoctoral fellow, Uhlenbeck and the Tinoco lab used biochemical methods to expand the database of sequences. Because WCF base pairing depends on strong, local hydrogen bonding and stacking interactions, a NN model was tested and found to fit the database. This suggested that the NN model would allow predictions for unmeasured sequences (105). In collaboration with related efforts in the Crothers lab, this led to original rules for predicting the thermodynamics of RNA folding (104).

Subsequent insights and research have continually improved success of the NN method. A rotational symmetry term was added to the model to account for the difference between duplexes formed by self- or non-self-complementary strands (42,74,107). Chemical synthesis on polymer supports allowed expansion of sequences available (41,42,108,109). Particularly important was addition of duplexes not beginning with multiple AU pairs and having melting temperatures near 37 °C, human body temperature. Analysis of the number of parameters allowed by the model (110,111) led to discovery that duplexes with the same nearest neighbors can have different thermodynamics depending on the terminal base pair (23). Applications of the method were expanded to larger RNAs by modifying dynamic programming algorithms to predict folding that optimized ΔG° (80,112,113) rather than base pairing (81). Recently, applications to even larger RNAs have become possible due to linearization of the algorithms (114,115).

NN parameters for canonical pairs have undergone substantial revisions over time, including treatment of end effects (23,24,26,42,105). GU terminal base pairs were initially assumed to be equivalent to AU terminal base pairs and were given the same penalty term (26). Expansion of the database and refitting of the model indicated that GU terminal base pairs do not require an end penalty (24). Measurements on helices with consecutive terminal GU pairs, however, revealed they are surprisingly more stable than predicted (25). Our updated NN model includes new parameters that account for end effects of both AU and GU pairs, including dependence on the penultimate pair.

Context-dependent variation of GU pair conformations (Figure 1) provides a structural rationale for treating terminal GU pairs differently. A fundamental assumption of a NN model is that strong local interactions dominate the energetic contributions determining conformation and stability for a particular nucleotide sequence. This implies that a stack of two WCF pairs will have the same thermodynamic stability in the middle of a helix as at the end of a helix. This NN approximation is consistent with structures of WCF RNA helices and the regular periodic shape of an RNA double helix. The diversity of GU pair conformations and stabilities, however, introduces variation into WCF paired helices. The unique structures of GU pairs facilitate binding recognition and specificity for metal ions, RNA tertiary interactions, protein interactions and drug binding (21). The challenge is incorporating this functionally important and structurally diverse motif into a NN model.

Prior models attempting to combine GU and WCF pairs into one set of thermodynamic parameters always had a few unexplained exceptions. For example, the motif 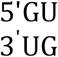 had one NN parameter value with an exception for the motif 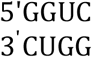, which had an extra bonus. Analysis of NMR structures and crystal structures of this motif, however, did not indicate a reason for this additional stability (51,52,116,117). In addition, the crystal structure of consecutive terminal GU pairs in (5’<u>GG</u>UGGCUG<u>UU</u>3’)_2_ had three slightly different helical conformations in the asymmetric unit but an overall remarkably A-form like structure that did not reveal a physical explanation for exceptional thermodynamic parameters (29). NMR studies of duplexes with consecutive terminal GU pairs usually showed broad resonances and few or weak NOEs in the final two GU pairs (25,29). Fluorescence and NMR studies have quantified different base pair dynamics in the middle and ends of helices for various types of base pairs (118-121). Consistent with this, the new NN model has increments for terminal nearest neighbors to distinguish them from internal nearest neighbors.

Interestingly, these increments are penalties +0.22 and +0.44 kcal/mol at 37 °C for an AU end pair on a penultimate AU or CG pair, respectively (Table 1). A similar penalty of +0.45 kcal/mol has previously been attributed to the presence of one fewer hydrogen bond when duplexes with identical nearest neighbors have two terminal AU pairs rather than terminal GC pairs (23). In contrast, incremental bonuses of -0.31 to -0.74 kcal/mol are assigned to terminal nearest neighbors consisting of an AU and a GU pair or two GU pairs. This would be consistent with a ΔS° bonus due to increased base pair dynamics at the ends of helices. For example, equal populations of three conformations at the end of a helix would provide a ΔG° bonus of -RT ln (3) = -0.68 kcal/mol at 37 °C.

The NN approximation is essential for efficient dynamic programming approaches to computing the minimum ΔG° secondary structure for an RNA sequence (80,81). In prior models, the special cases for GU pairs required additional considerations in dynamic programming algorithm computations. A helix end occurs not only at the 5’ and 3’ ends of an RNA molecule but also at every junction, internal loop, hairpin loop, and mismatch or bulge in an RNA secondary structure. GU helix end pairs occur at in accepted RNA secondary structures at a rate of approximately 13 per 1000 bases in the sequence (Supplemental Figure 11). They occur at a much higher rate in predicted structural ensembles. Thus, consideration of GU pairs and special rules for positional dependence present a frequent step in the computations. The NN parameter model presented here improves predictions for sequences and structures with terminal AU and GU pairs and will also accelerate computation of the minimum free energy structure for any sequence.

For example, several terminal AU and GU motifs occur in the secondary structure for the Ψ packaging sequence in HIV-1 RNA (20) and the motif 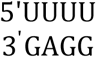 binds a novel drug. Each helix in the three-way junction that binds the drug has an AU or GU pair at the end, and the new NN parameters in this work would estimate that the 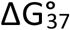 for these three helices is at least 0.9 kcal/mol more stable than current predictions.

Another recent example is the SL3 helix that forms between the 5’ and 3’ ends of SARS-CoV-2. This helix has been identified experimentally (122) and computationally (123). One end of SL3 terminates in a 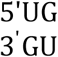 nearest neighbor. Results in Table 1B assign a 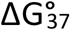 of -0.38 -0.74 = -1.12 kcal/mol to this end, which is more stable than previous predictions.

Free energy predictions from nearest neighbors for RNAs secondary structures provide the base line for analysis of the stabilities of RNA interactions with drugs and proteins, and thus provide a foundational resource for RNA structure and function studies. Our future analyses will evaluate the impact of the new NN parameters on the thermodynamic parameters for mismatches, internal loops, bulges, and helix junctions. These loop motifs form many of the recognition sites for proteins, metal ions, and therapeutics.

To illustrate a NN calculation to estimate helix stability, Figure 4 provides two example calculations. The first is the sequence (5’UGUCGAUA)_2_, with experimental stability provided in Table 2. The second is 5’UAGGUCAG paired with 5’CUGGUCUA. This calculation illustrates that the 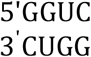 motif, an outlier in prior nearest neighbor models, is now handled with nearest neighbor stacks. An Excel spreadsheet is provided with the Supplementary Materials to calculate user-inputted helical NN stabilities.

**Figure 4.**
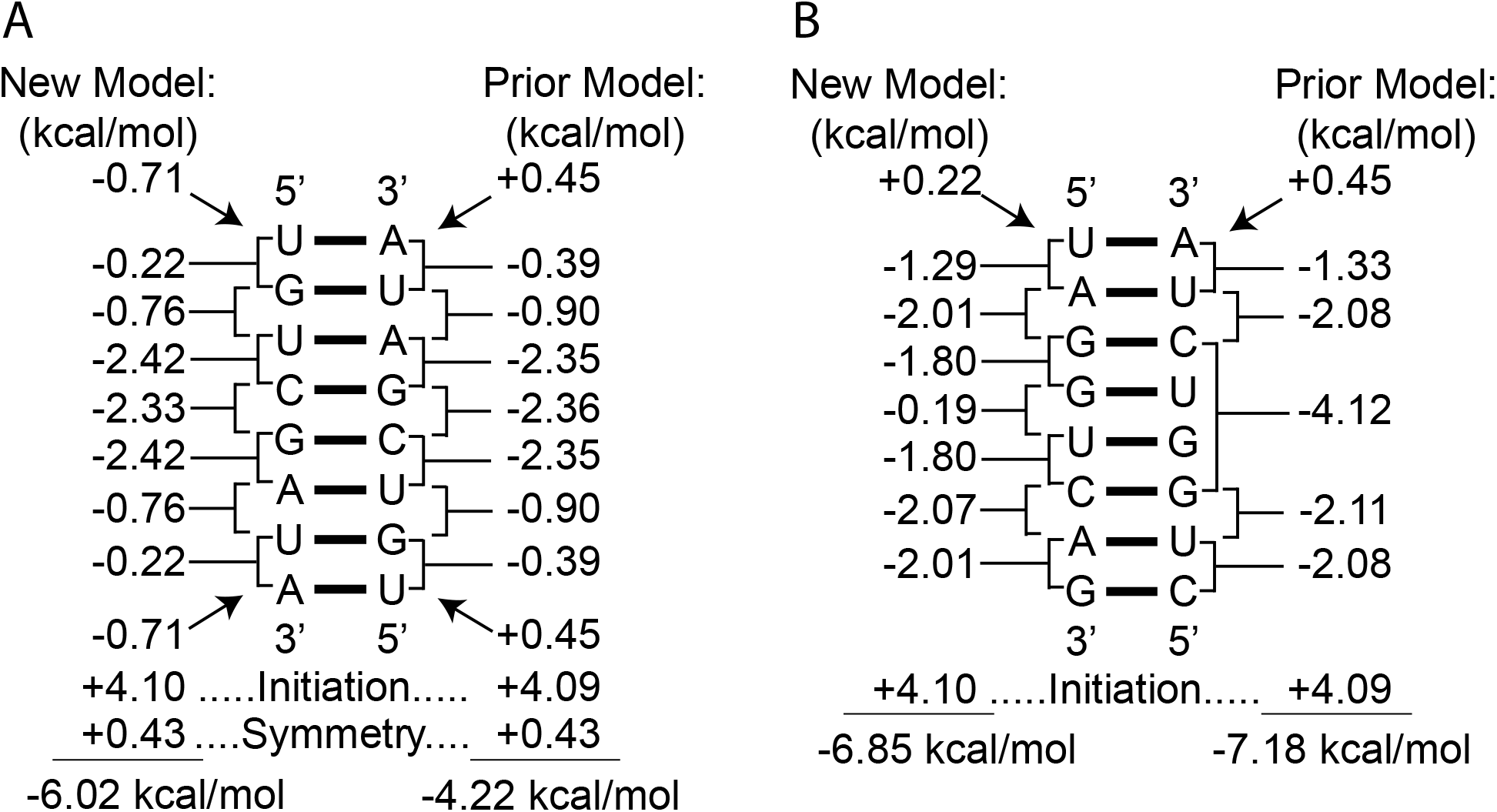
Example calculations of helical 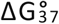. Panel A shows the stability calculation for (5’UGUCGAUA)_2_, which is shown in Table 2 to have an experimentally determined 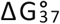 of -6.10 kcal/mol. This sequence is self-complementary and therefore the symmetry penalty is added. Panel B shows the stability calculation for 5’UAGGUCAG paired to 5’CUGGUCUA. This demonstrates the difference in treatment for the (GGUC)_2_ motif. For both sequences, calculations are provided for the current parameters derived here and the previous parameters (23,24). The total stability is the sum of the stability increments.

In summary, the updated NN model is consistent with previous parameters for WCF pairs, includes new parameters accounting for increased base pair dynamics at ends for helices ending in AU or GU pairs, improves predictions for duplexes with terminal AU or GU pairs, and resolves a prior exceptional parameter for a specific GU motif. The model for the NN parameters has low uncertainty in ΔG° and ΔH° and low correlations between parameters. The statistically robust model maintains the physical basis that differences in hydrogen bonding, stacking, and nucleotide dynamics determine the sequence dependence of NN base stacks. The new thermodynamic parameters will help improve RNA structure prediction tools and facilitate discoveries in RNA biology, catalysis, and therapeutics.

## Supporting information

Worksheet for estimating duplex stabilities.

Database of data used for linear regression.

## Supplemental Information to Accompany

**Supplemental Table 1.**
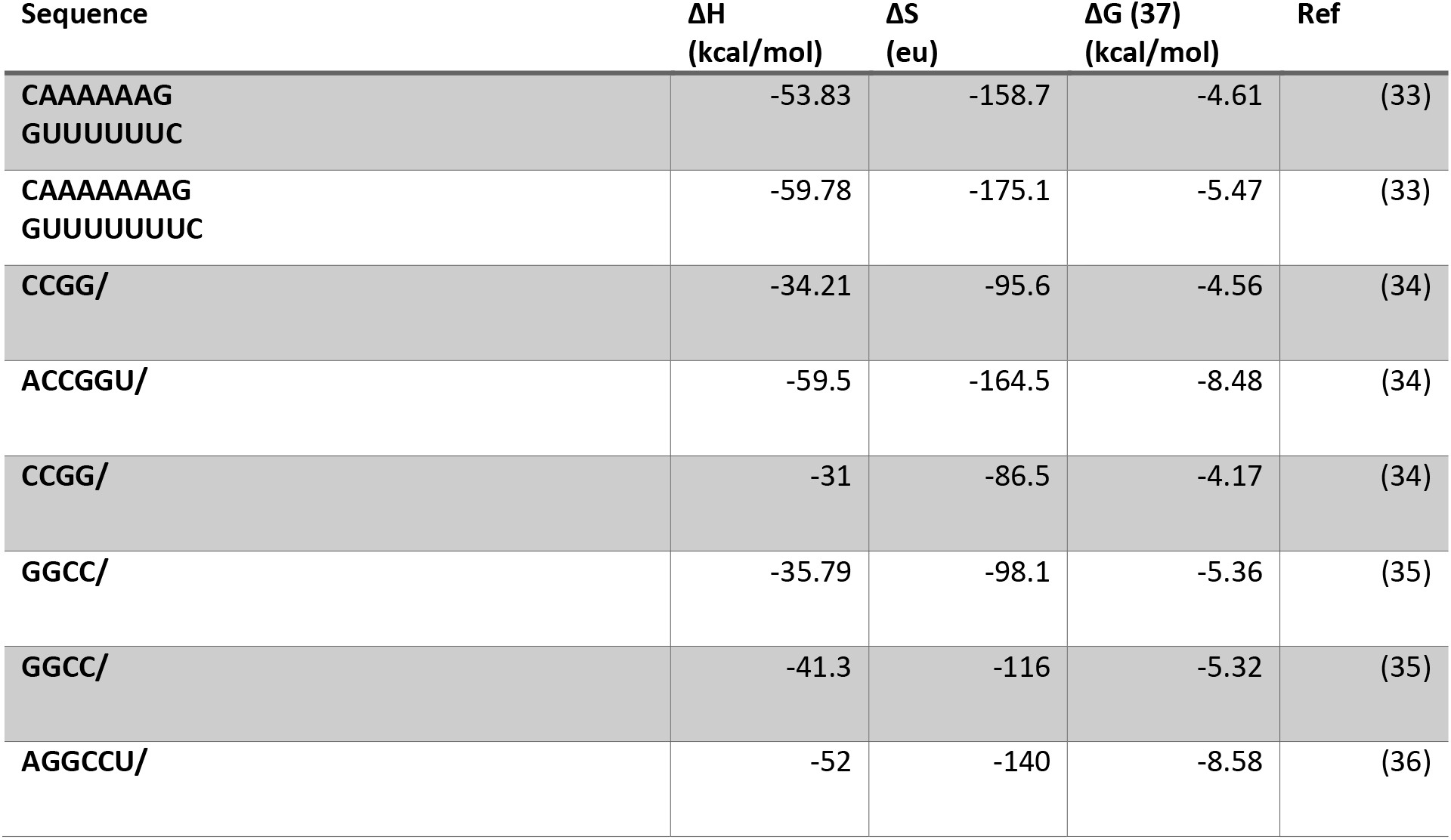
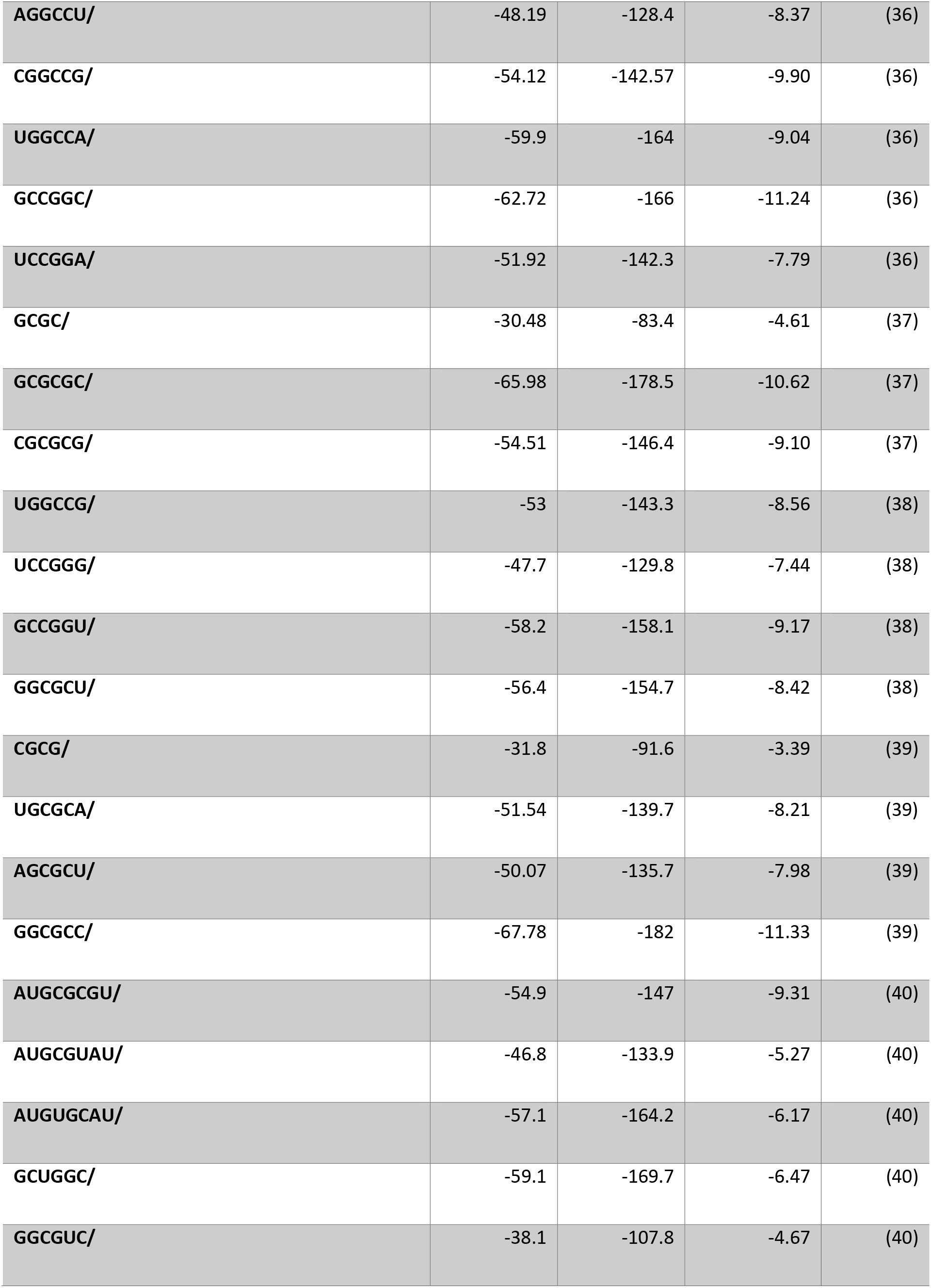
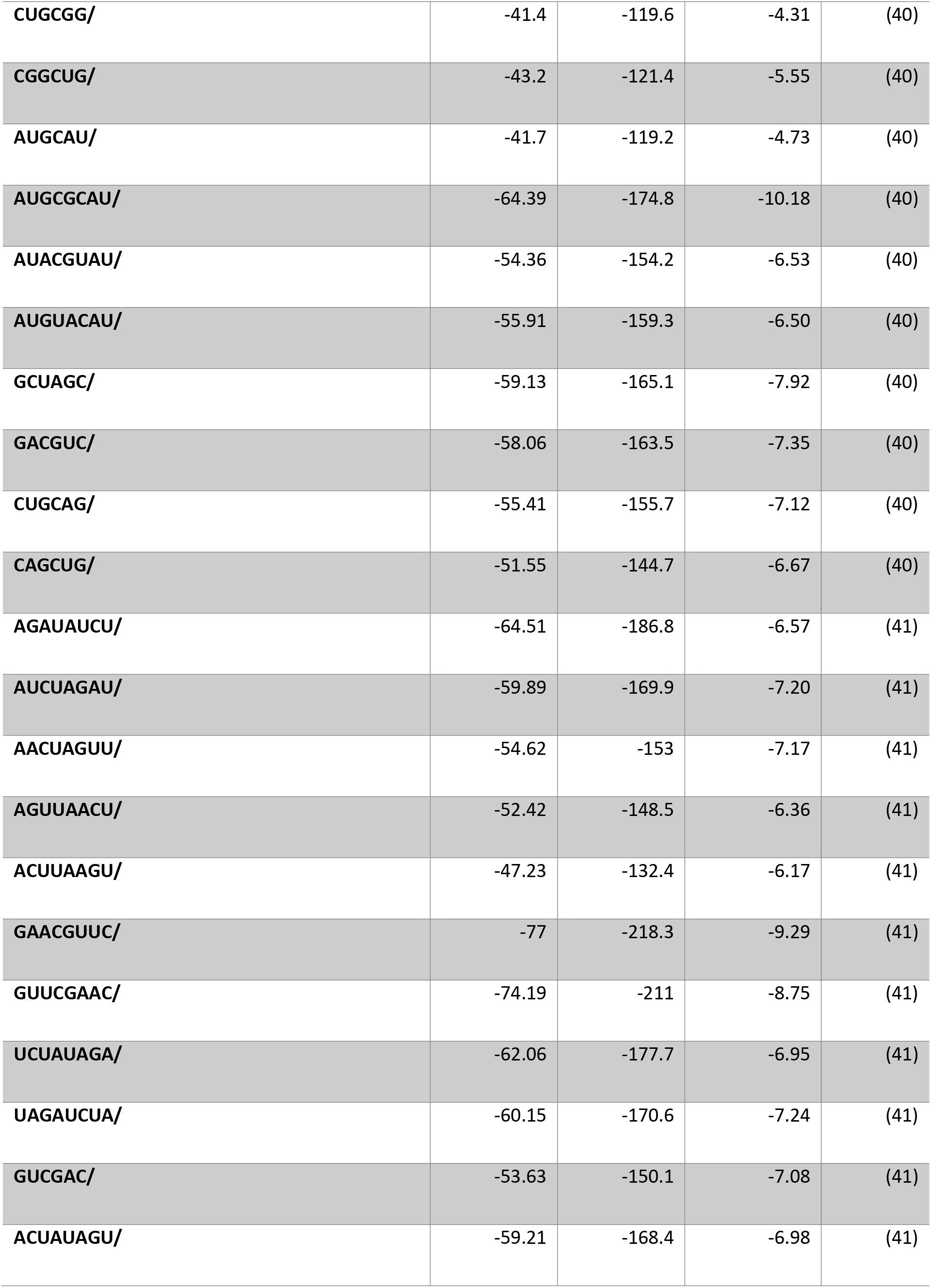
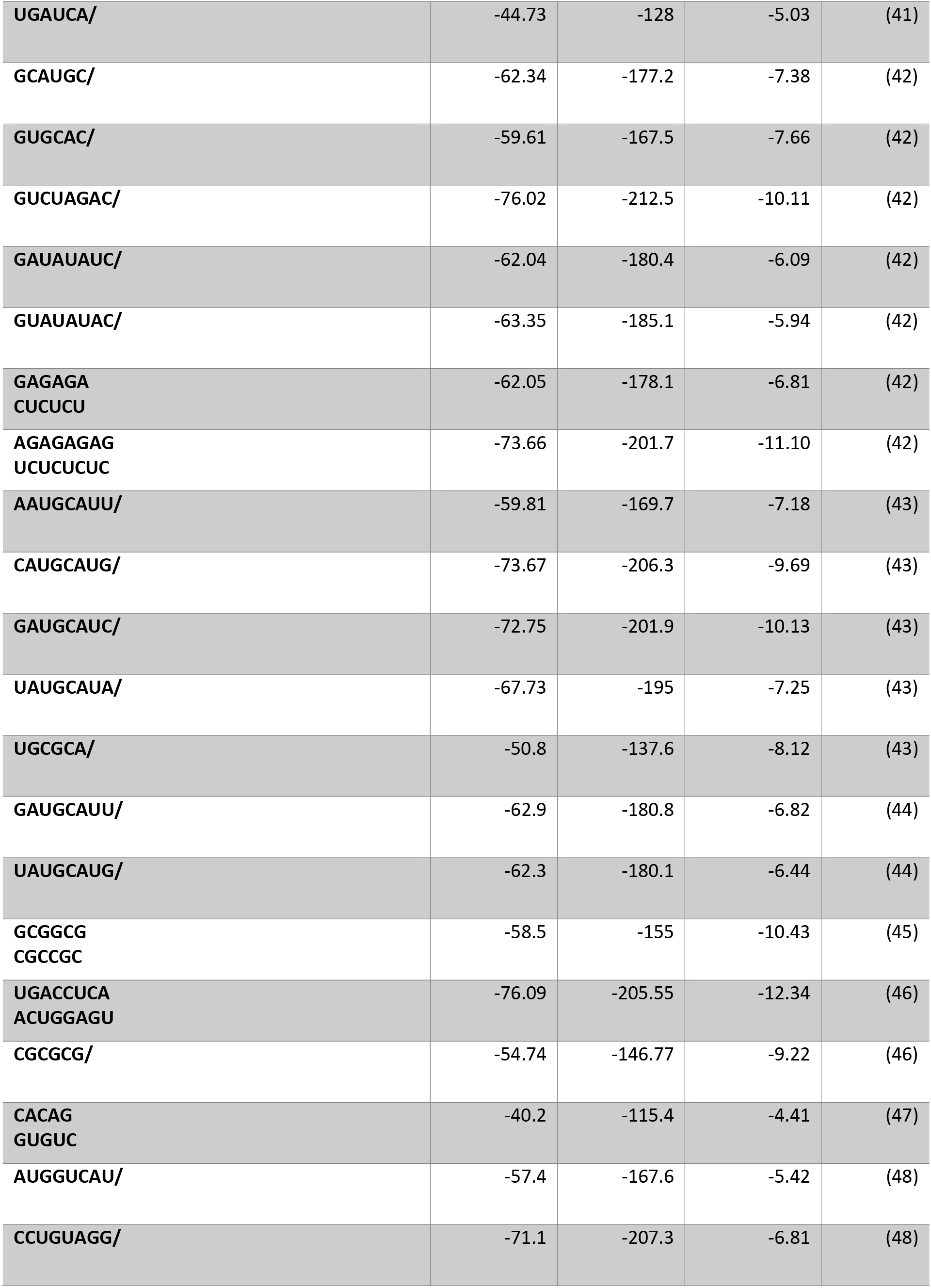
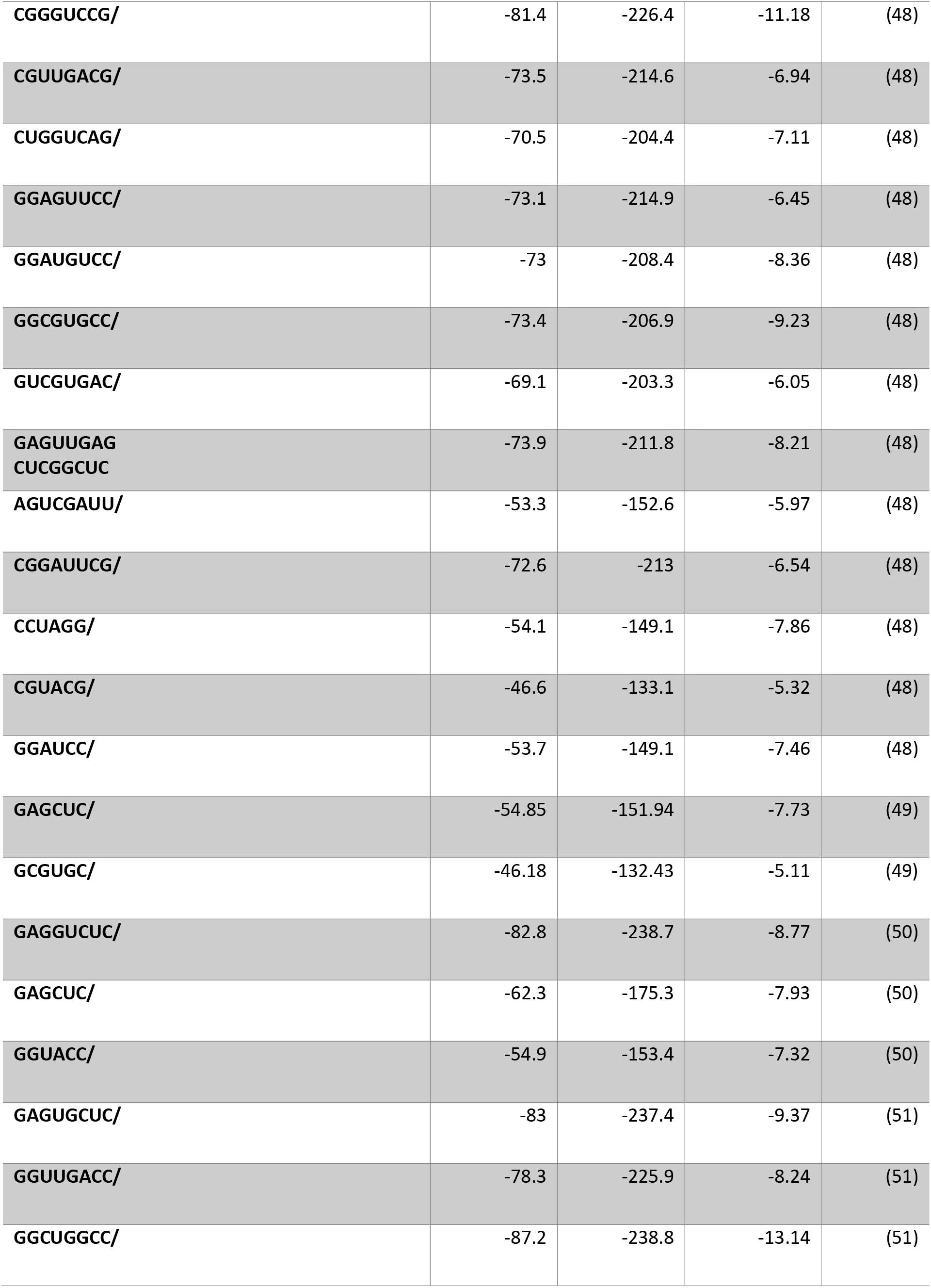
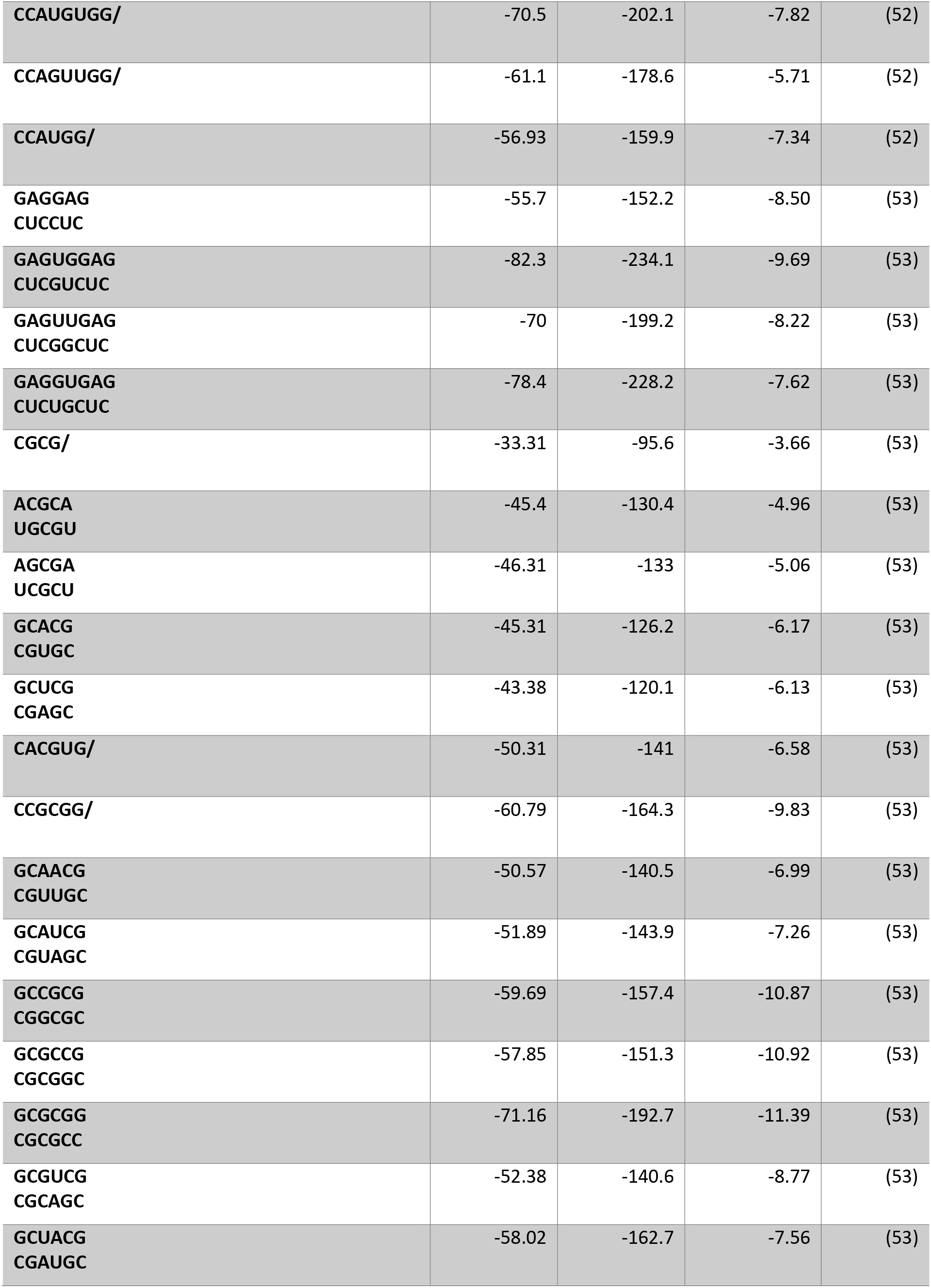
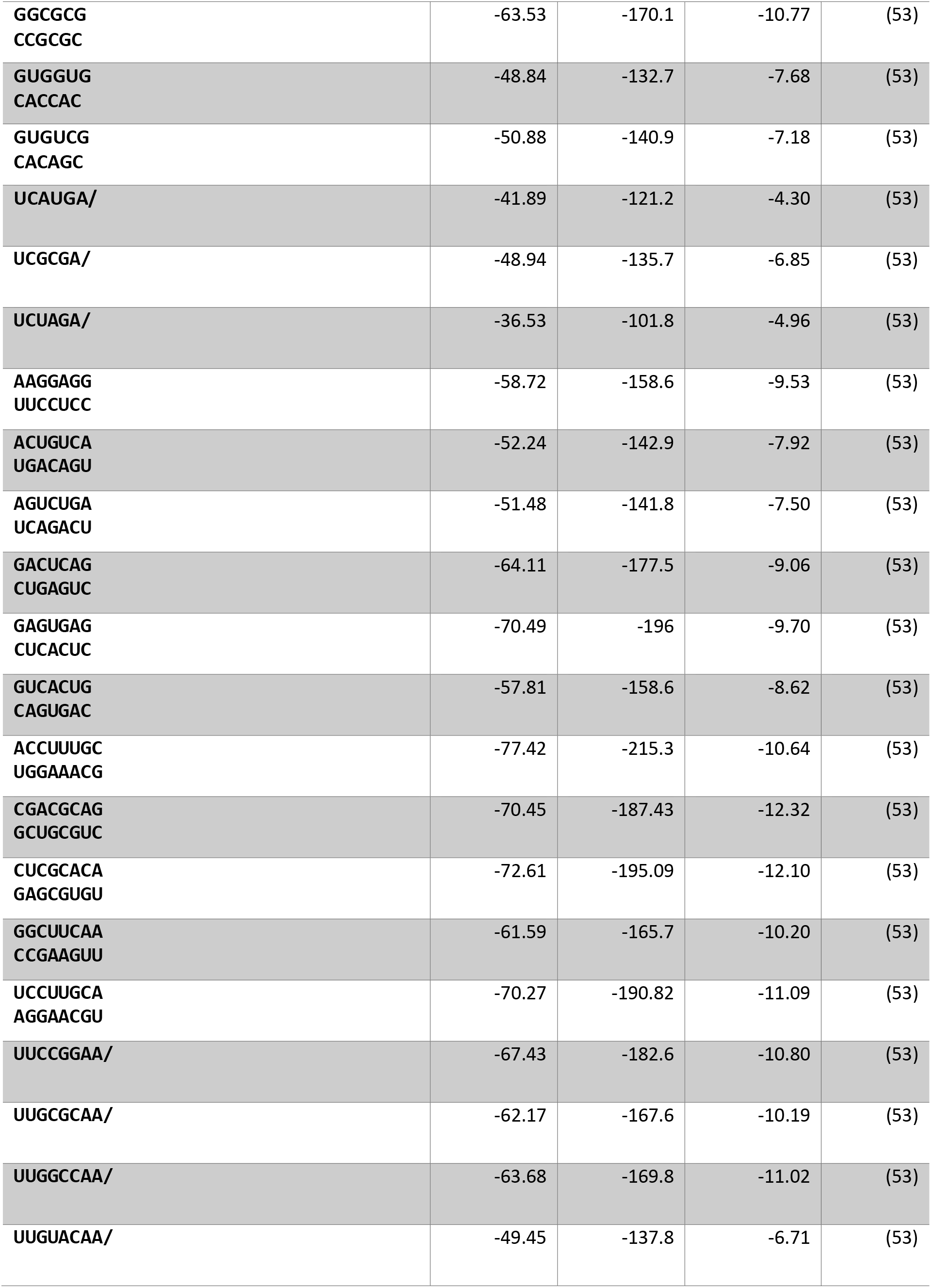
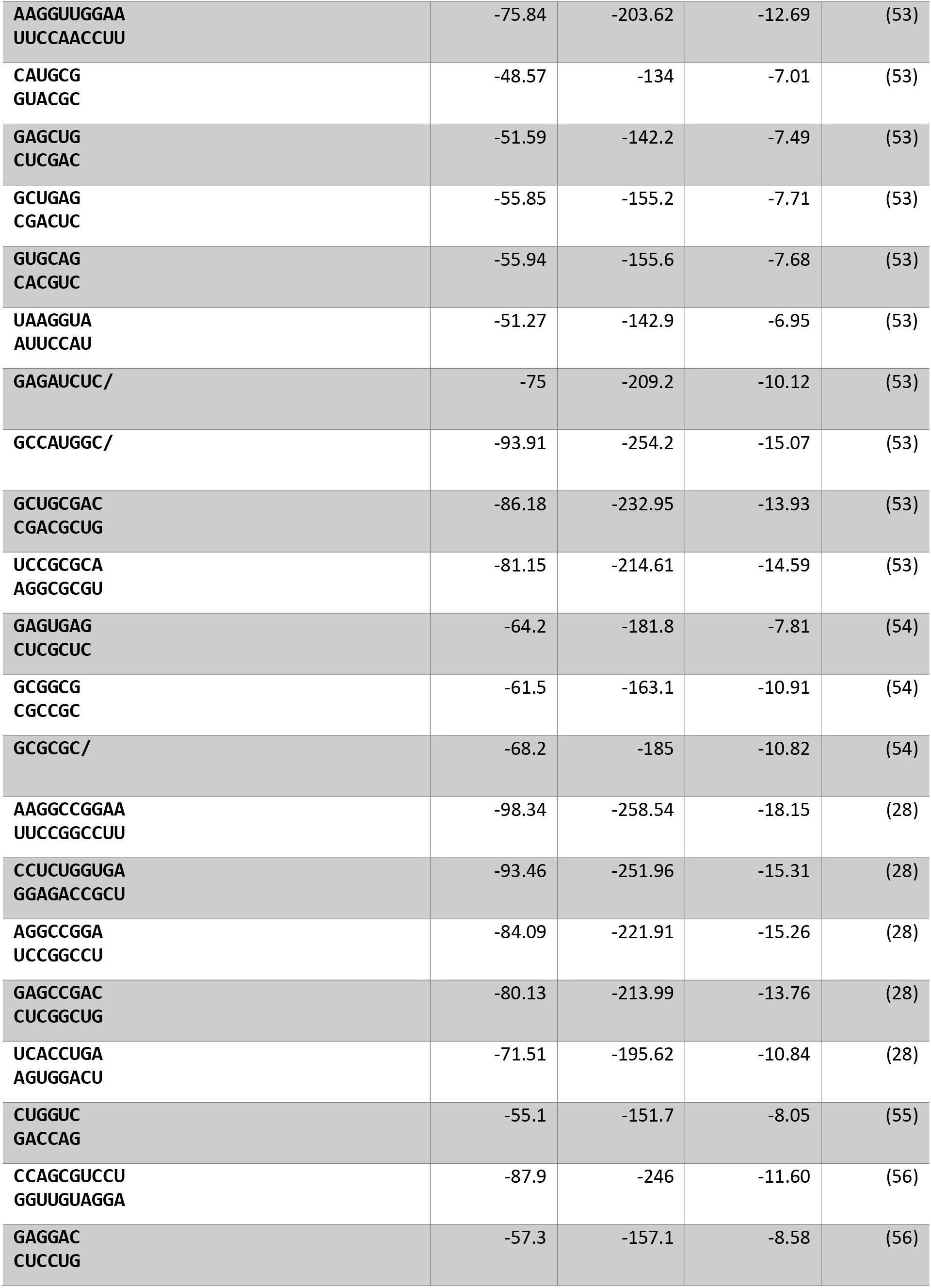
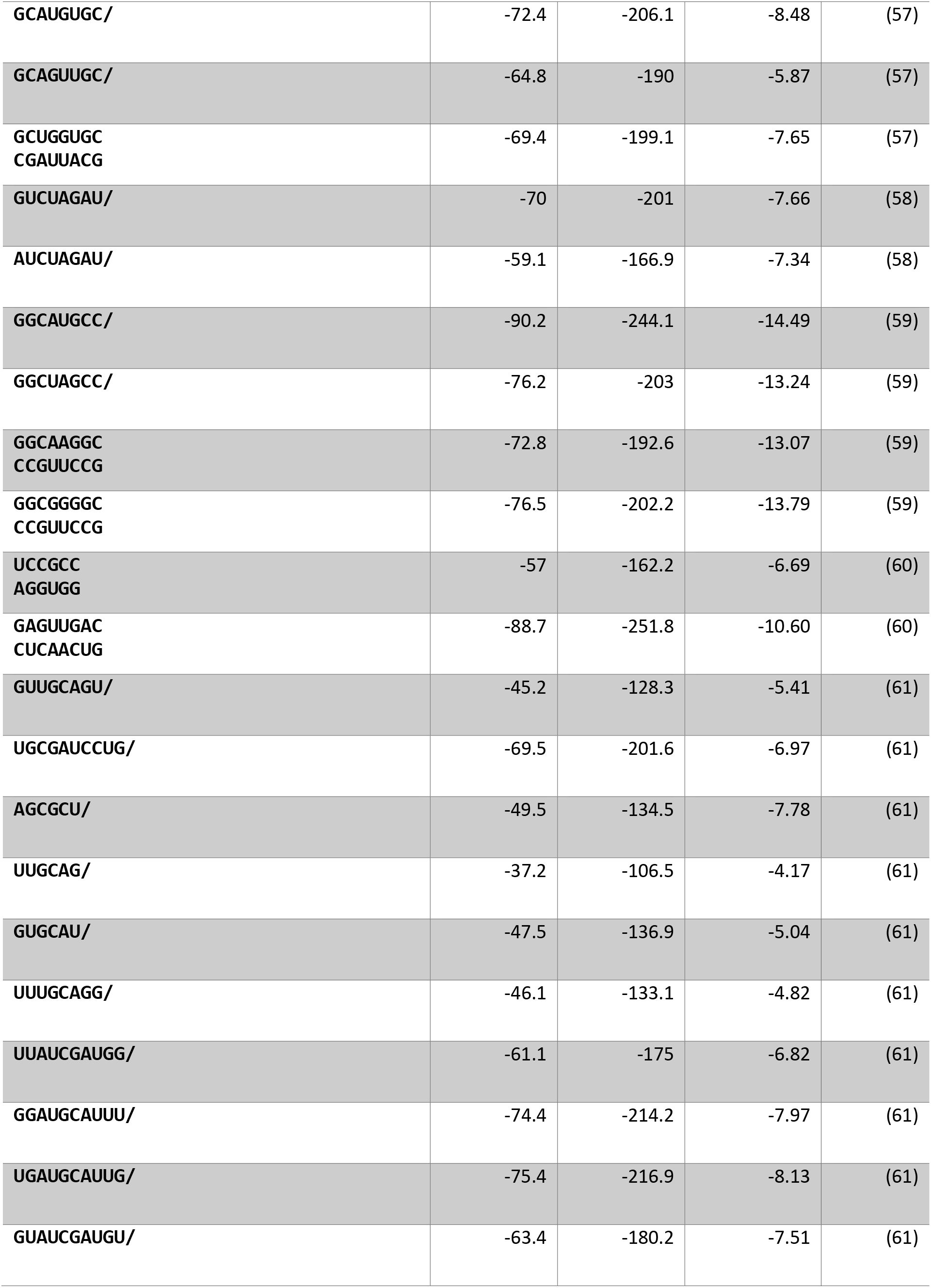
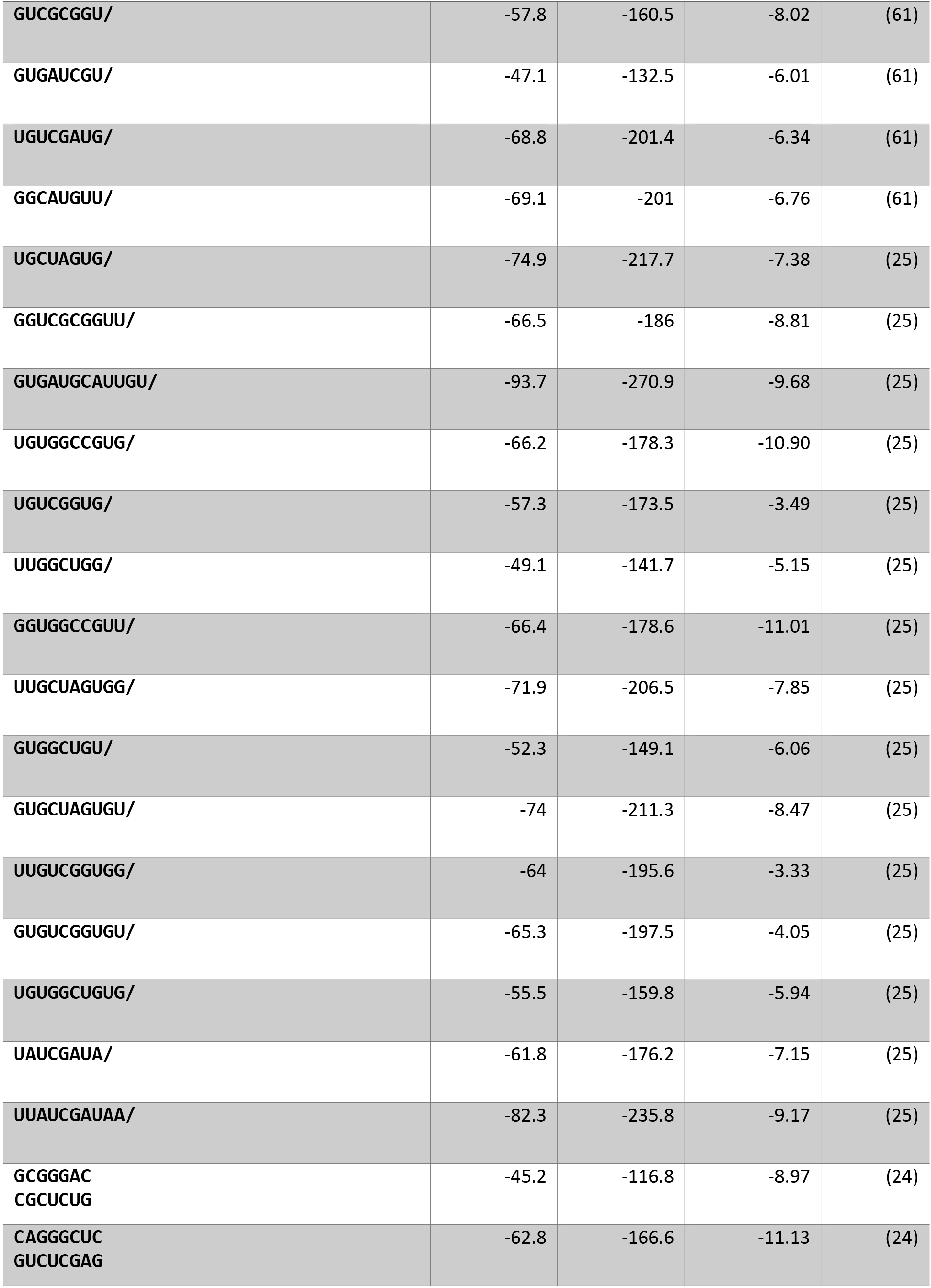
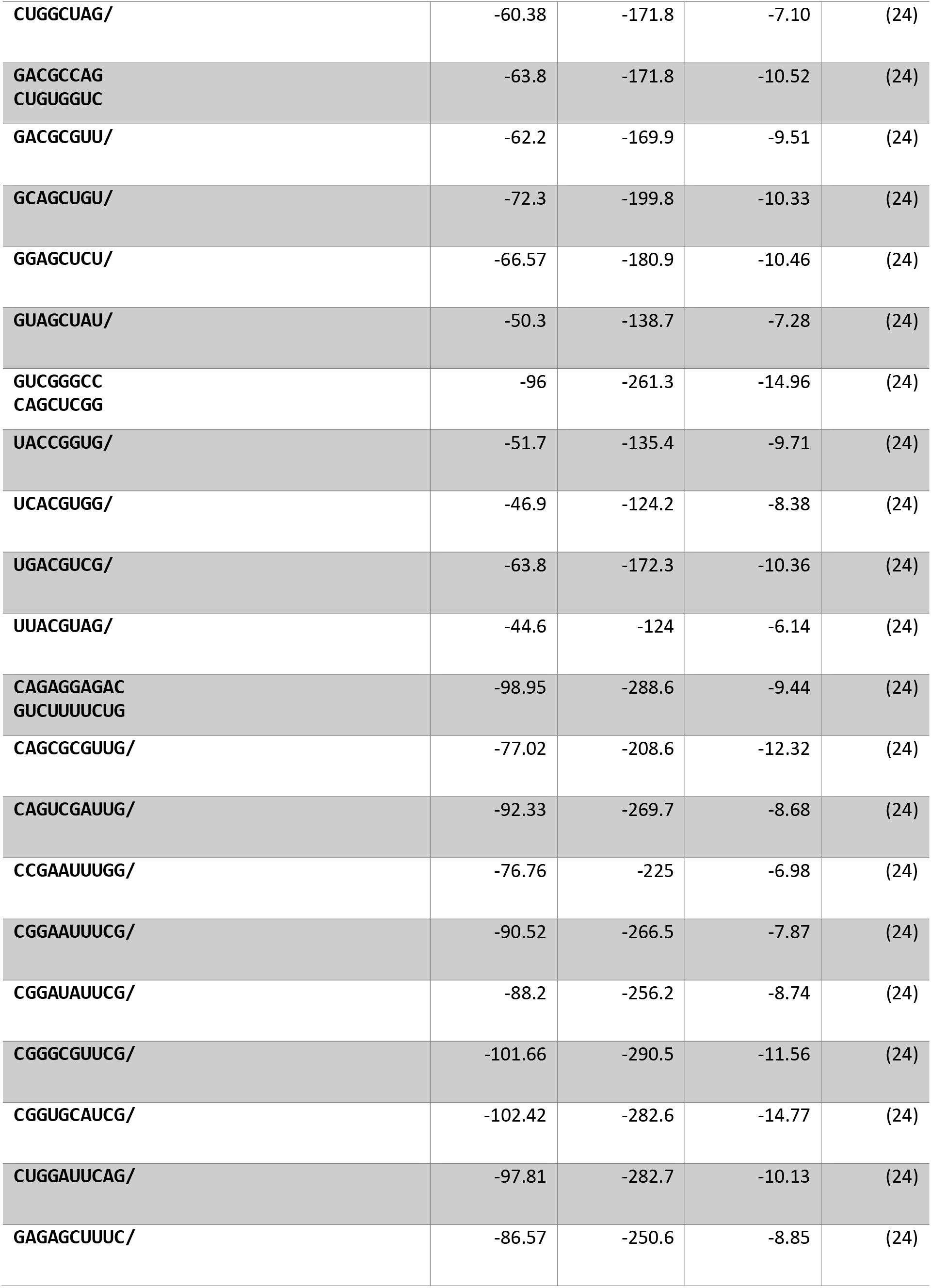
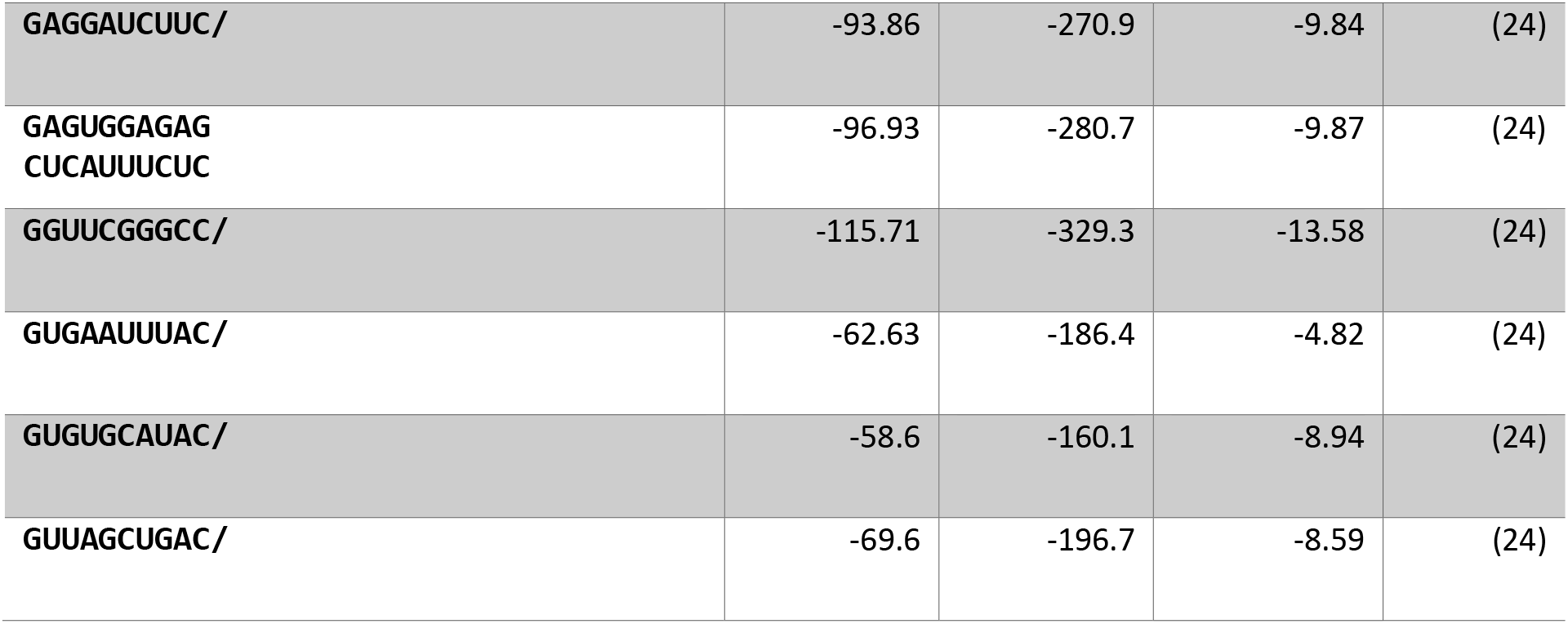
Duplex sequences used for parameter fitting.

**Supplemental Table 2.** A) WCF helical parameters annotated with errors estimated as standard errors of regression. These error estimates are not as good as those in Table 1 derived by perturbing the input experimental data. B) GU stack parameters annotated with errors estimated as standard errors of regression. These error estimates are not as good as those in Table 1 derived by perturbing the input experimental data.

| Feature | New Model |  | 1998 Model <sup>†</sup> |  |
| --- | --- | --- | --- | --- |
| | $\Delta G_{37}^{\circ}$<br>(kcal/mol) | $\Delta H$<br>(kcal/mol) | $\Delta G_{37}^{\circ}$<br>(kcal/mol) | $\Delta H$<br>(kcal/mol) |
| GC/CG | -3.46 ± 0.07 | -16.52 ± 0.84 | -3.42 ± 0.08 | -14.88 ± 1.58 |
| CC/GG | -3.28 ± 0.05 | -13.94 ± 0.66 | -3.26 ± 0.07 | -13.39 ± 1.24 |
| GA/CU | -2.42 ± 0.05 | -13.75 ± 0.64 | -2.35 ± 0.06 | -12.44 ± 1.20 |
| CG/GC | -2.33 ± 0.08 | -9.61 ± 1.00 | -2.36 ± 0.09 | -10.64 ± 1.65 |
| AC/UG | -2.25 ± 0.06 | -11.98 ± 0.76 | -2.24 ± 0.06 | -11.40 ± 1.23 |
| CA/GU | -2.07 ± 0.06 | -10.47 ± 0.76 | -2.11 ± 0.07 | -10.44 ± 1.28 |
| AG/UC | -2.01 ± 0.06 | -9.34 ± 0.78 | -2.08 ± 0.06 | -10.48 ± 1.24 |
| UA/AU | -1.29 ± 0.09 | -9.16 ± 1.10 | -1.33 ± 0.09 | -7.69 ± 2.02 |
| AU/UA | -1.09 ± 0.08 | -8.91 ± 1.01 | -1.10 ± 0.08 | -9.38 ± 1.68 |
| AA/UU | -0.94 ± 0.04 | -7.44 ± 0.52 | -0.93 ± 0.03 | -6.82 ± 0.79 |
| Initiation | +4.10 ± 0.20 | +4.66 ± 2.47 | +4.09 ± 0.22 | +3.61 ± 4.12 |
| AU End on AU | +0.22 ± 0.06 | +4.36 ± 0.76 | +0.45 ± 0.04 <sup>‡</sup> | +3.72 ± 0.83 <sup>‡</sup> |
| AU End on CG | +0.44 ± 0.04 | +3.17 ± 0.55 | +0.45 ± 0.04 <sup>‡</sup> | +3.72 ± 0.83 <sup>‡</sup> |

| Feature | New Model |  | 2012 Model <sup>¶</sup> |  |
| --- | --- | --- | --- | --- |
| | $\Delta G_{37}^{\circ}$<br>(kcal/mol) | $\Delta H$<br>(kcal/mol) | $\Delta G_{37}^{\circ}$<br>(kcal/mol) | $\Delta H$<br>(kcal/mol) |
| GC/UG | -2.23 ± 0.12 | -14.73 ± 1.32 | -2.15 ± 0.10 | -11.09 ± 1.78 |
| CU/GG | -1.93 ± 0.13 | -9.26 ± 1.45 | -1.77 ± 0.09 | -9.44 ± 1.76 |
| GG/CU | -1.80 ± 0.11 | -12.41 ± 1.28 | -1.80 ± 0.09 | -7.03 ± 1.75 |
| CG/GU | -1.05 ± 0.12 | -5.64 ± 1.37 | -1.25 ± 0.09 | -5.56 ± 1.68 |
| AU/UG | -0.76 ± 0.13 | -9.23 ± 1.50 | -0.90 ± 0.08 | -7.39 ± 1.65 |
| GA/UU | -0.60 ± 0.11 | -10.58 ± 1.26 | -0.51 ± 0.08 | -10.38 ± 1.79 |
| UG/GU | -0.38 ± 0.15 | -8.76 ± 1.75 | -0.57 ± 0.19 | -12.64 ± 4.01 |
| UA/GU | -0.22 ± 0.13 | -2.72 ± 1.47 | -0.39 ± 0.09 | -0.96 ± 1.80 |
| GG/UU | -0.20 ± 0.15 | -9.06 ± 1.71 | -0.25 ± 0.16 | -17.82 ± 3.75 |
| GU/UG | -0.19 ± 0.16 | -7.66 ± 1.85 | +0.72 ± 0.19 | -13.83 ± 4.21 |
| AG/UU | +0.02 ± 0.12 | -5.10 ± 1.38 | -0.35 ± 0.08 | -3.96 ± 1.73 |
| GGUC/CUGG | (-3.79 ± 0.27) <sup>†</sup> | (-32.48 ± 3.16) <sup>†</sup> | -4.12 ± 0.54 | -30.80 ± 8.87 |
| AU End on GU | -0.71 ± 0.30 | 5.16 ± 3.42 | +0.45 ± 0.04 <sup>*</sup> | +3.72 ± 0.83 <sup>*</sup> |
| GU End on CG | 0.13 ± 0.14 | 3.91 ± 1.63 | 0.00 ± 0.00 <sup>‡</sup> | 0.00 ± 0.00 <sup>‡</sup> |
| GU End on AU | -0.31 ± 0.14 | 3.65 ± 1.56 | 0.00 ± 0.00 <sup>‡</sup> | 0.00 ± 0.00 <sup>‡</sup> |
| GU End on GU | -0.74 ± 0.16 | 6.23 ± 1.89 | 0.00 ± 0.00 <sup>‡</sup> | 0.00 ± 0.00 <sup>‡</sup> |

**Supplemental Table 3:**
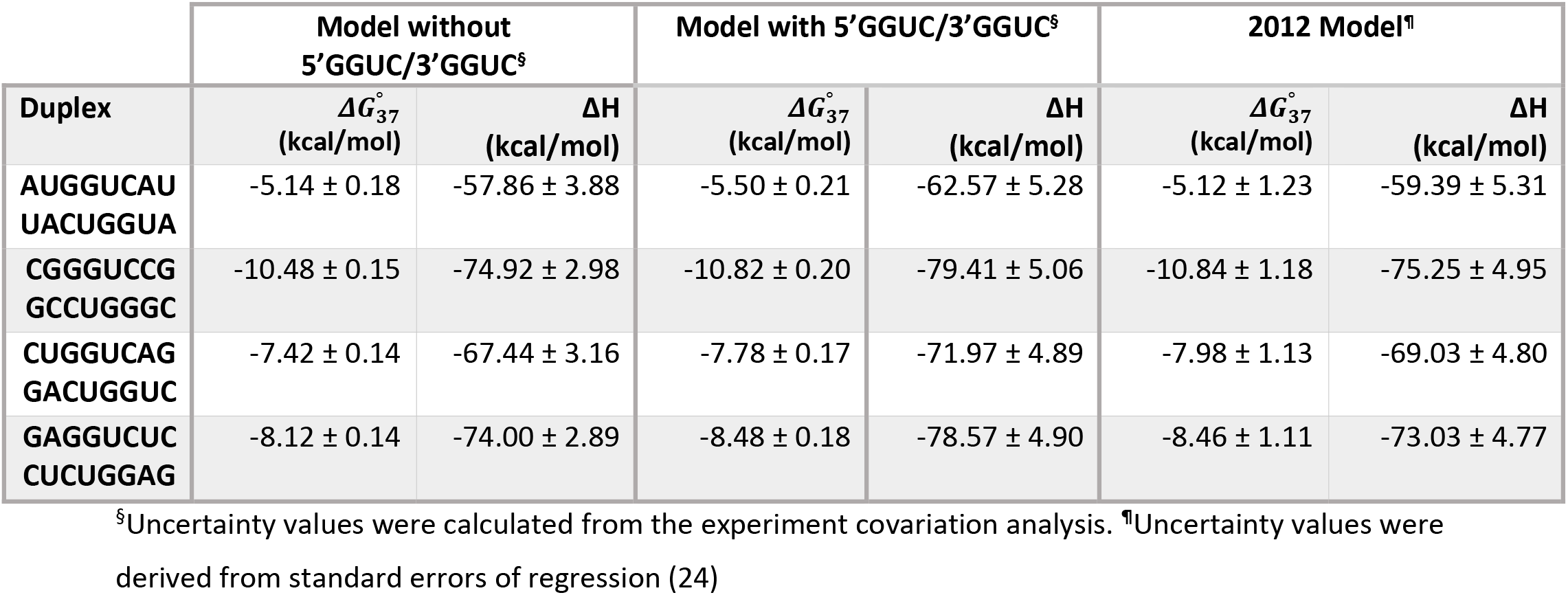
Comparison of predictions of models with and without 5’GGUC/3’CUGG parameter.

**Supplemental Figure 1:**
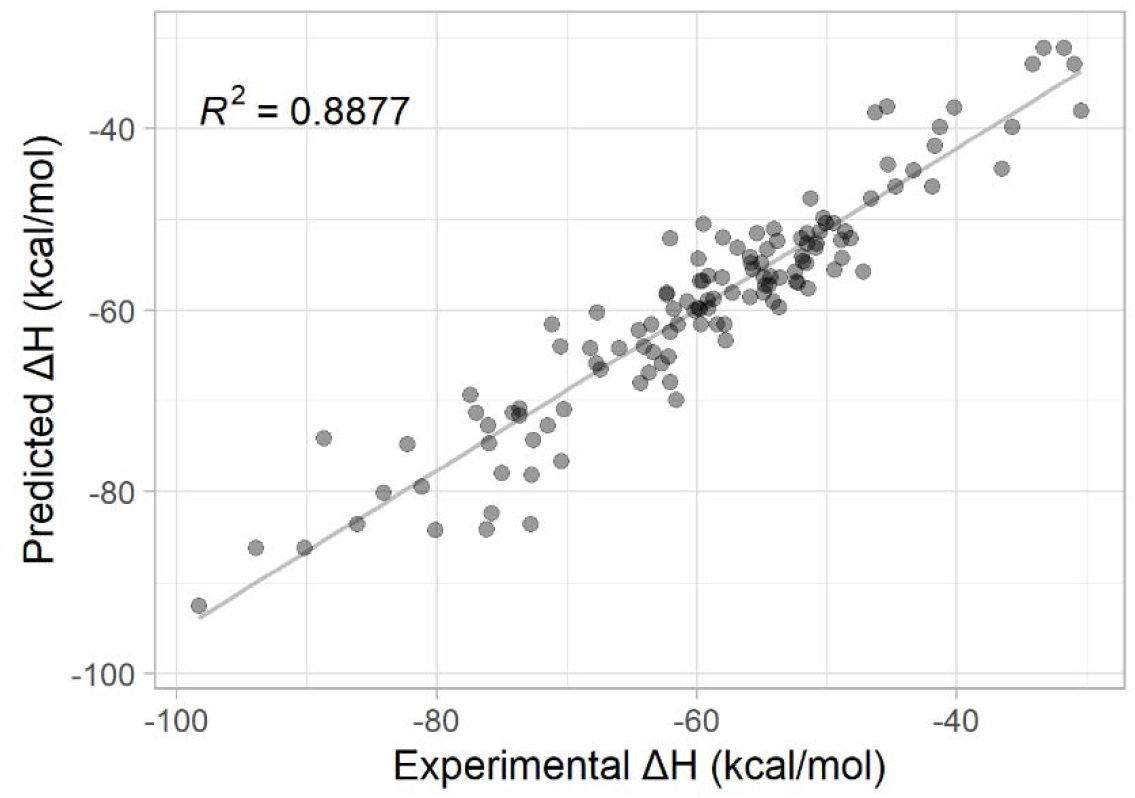
ΔH° predictions as a function of experimentally-determine ΔH° for canonical WCF nearest neighbor parameters. ΔH° values predicted from updated nearest neighbor parameters for duplexes composed solely canonical WCF base pairs in Table 1A plotted against values determined from optical melting experiments.

**Supplemental Figure 2:**
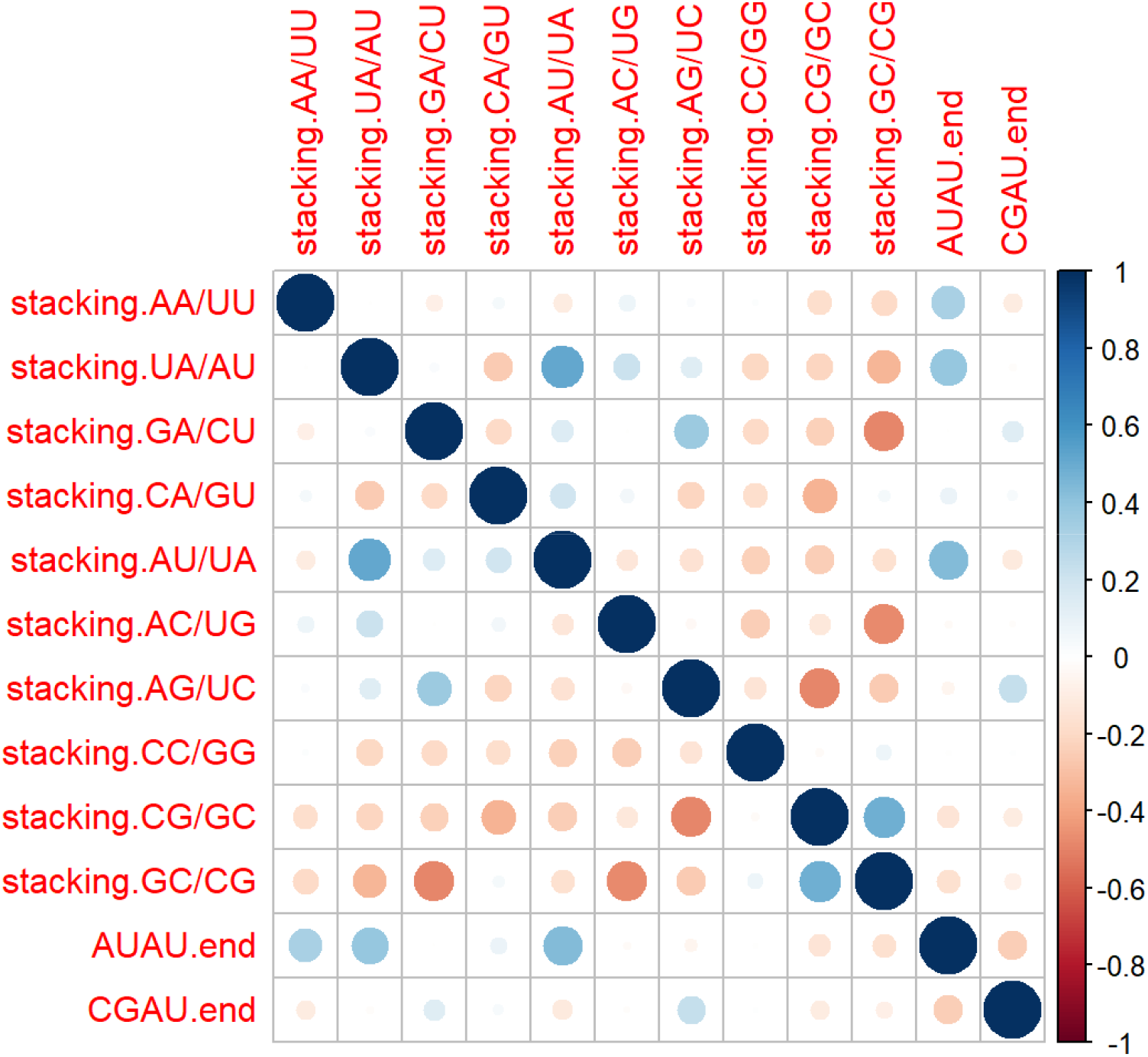
Pearson correlations between model features. Pairwise Pearson correlation coefficients were calculated for the observed frequencies of WCF stacking parameters. The correlation coefficients are encoded in the size and color of each circle.

**Supplemental Figure 3:**
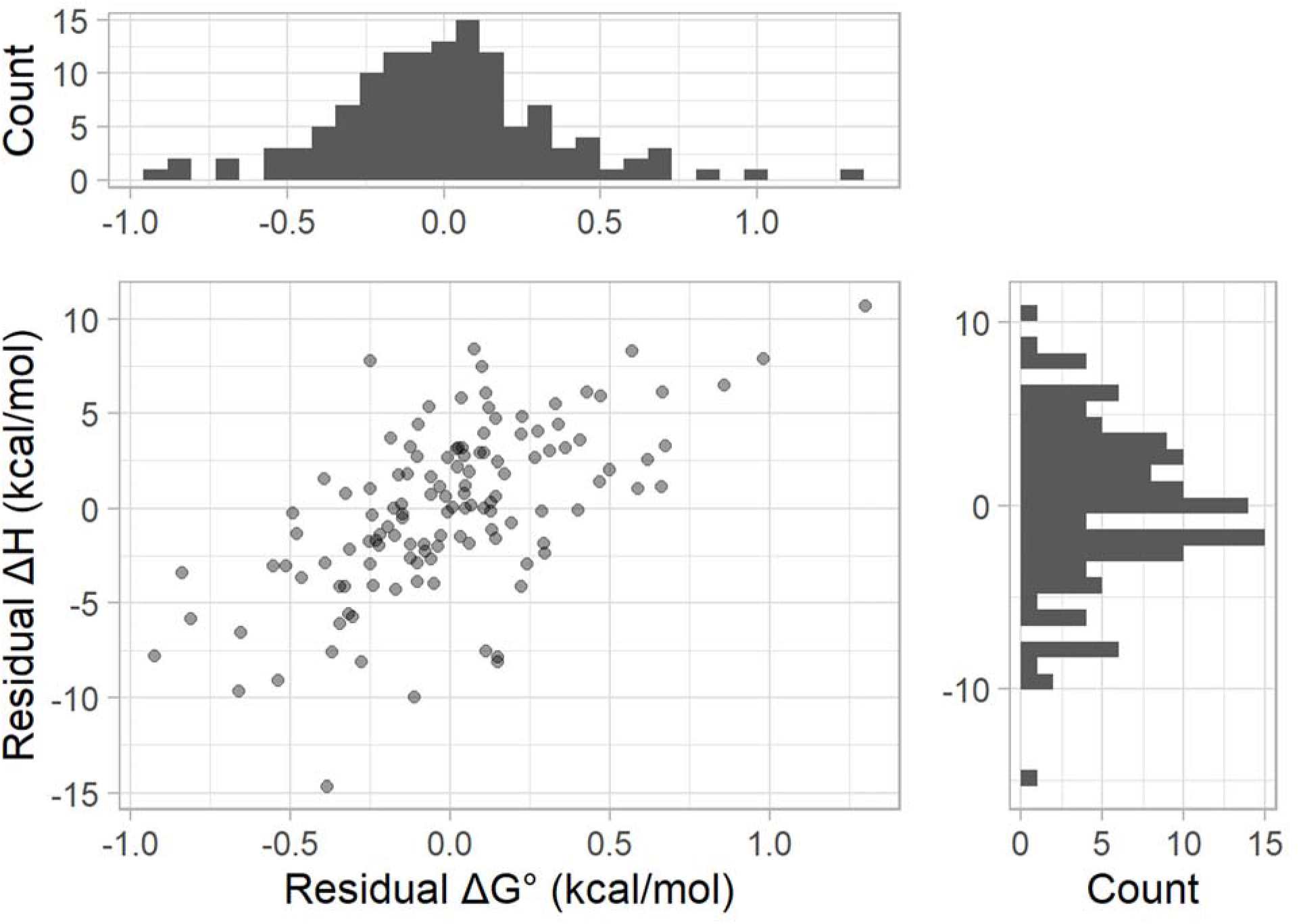
Residuals for the Watson-Crick-Franklin stacking parameter predictions. Shown are the scatter plot between the residuals in ΔH° and folding ΔG°_37_ change predictions (main plot), a histogram of the ΔH° residuals (right plot), and a histogram of the ΔG°_37_. residuals (top plot).

**Supplemental Figure 4:**
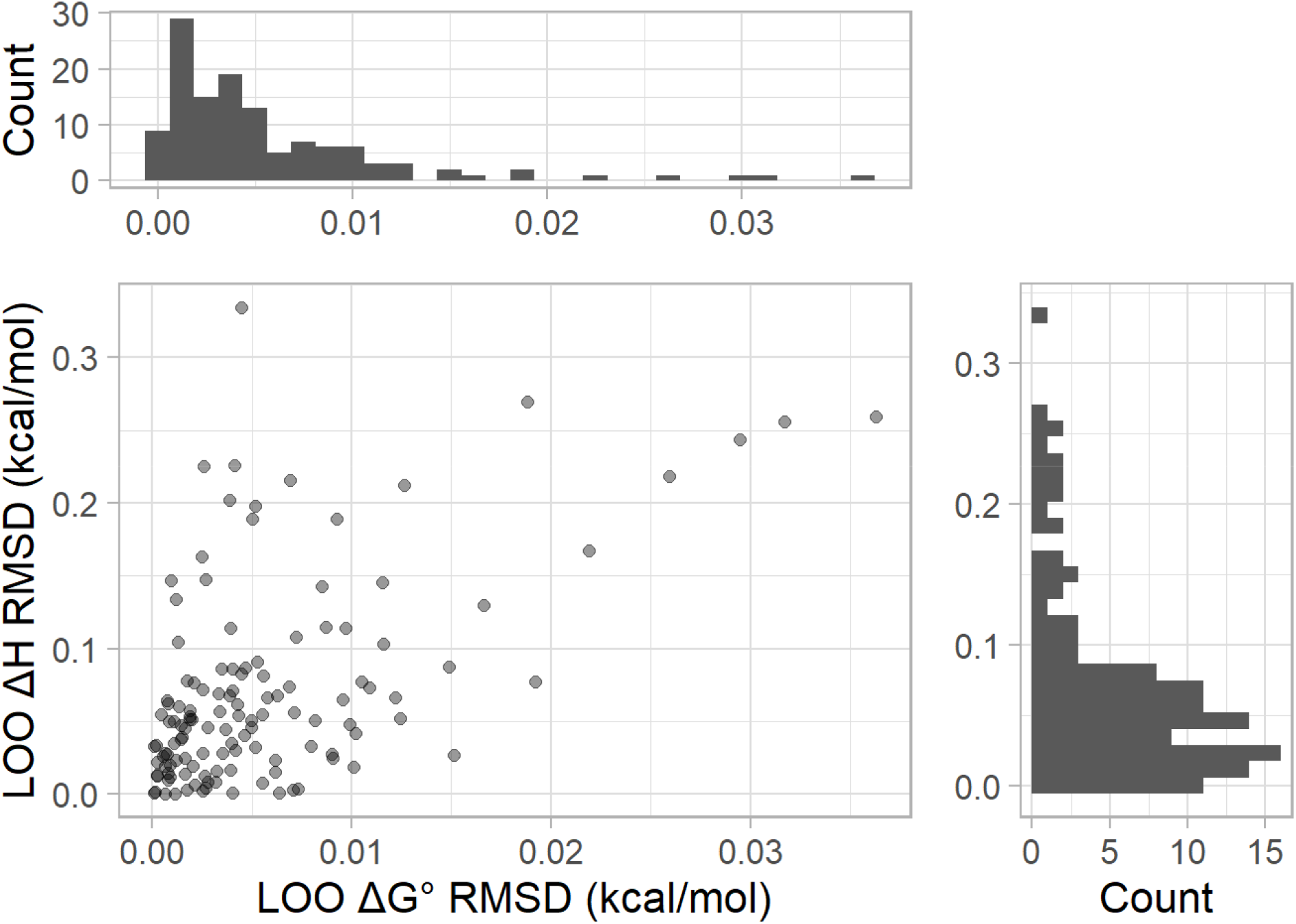
Leave-One-Out (LOO) analysis for WCF helix parameters. The plotted value is the RMSD of the parameter values for leaving out a single experiment as compared to when all experiments are used.

**Supplemental Figure 5:**
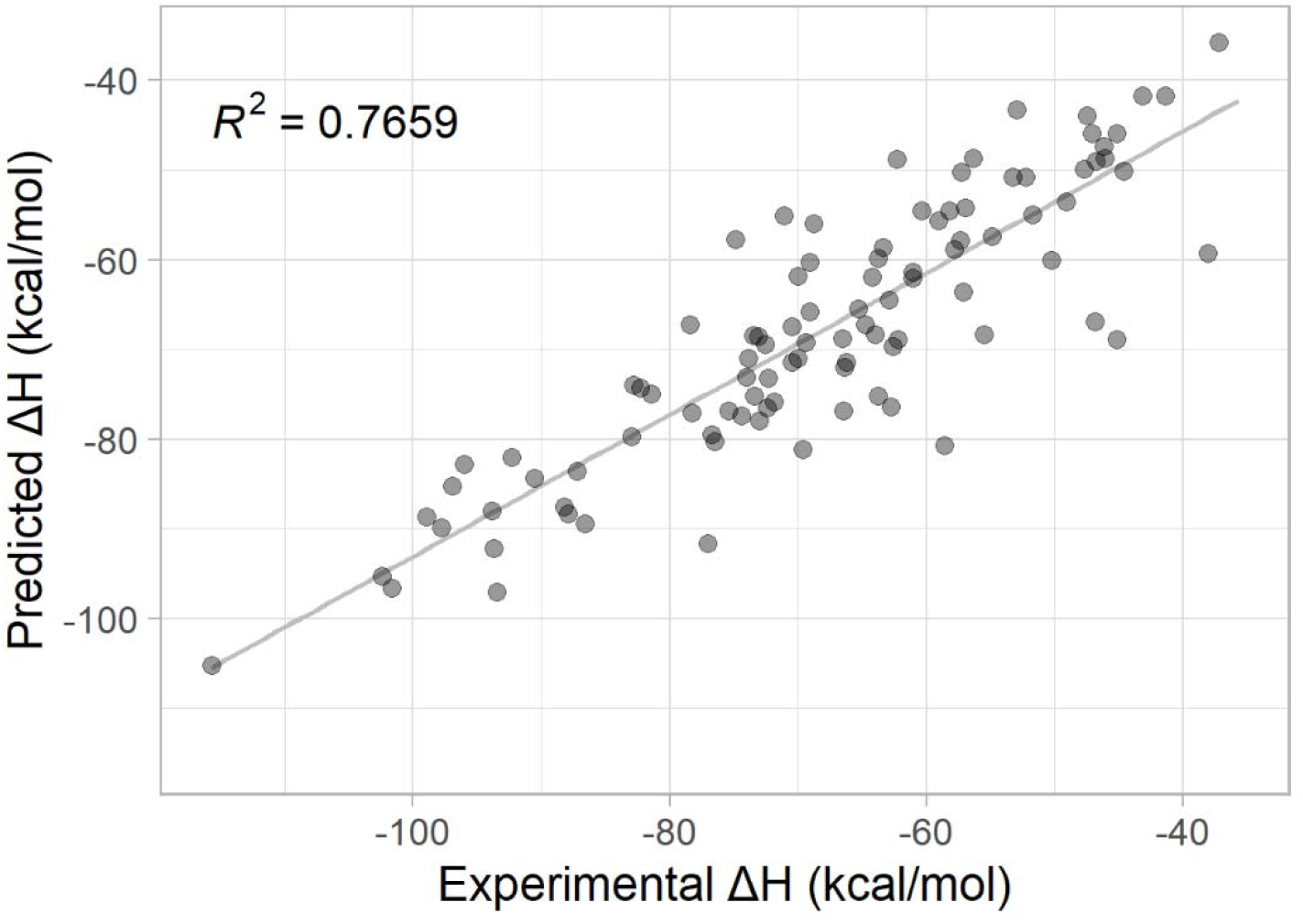
Correlation between predicted and observed ΔH° for duplexes with WCF and GU pairs. ΔH° values predicted from parameters in Table 1 plotted against values determined from optical melting experiments.

**Supplemental Figure 6:**
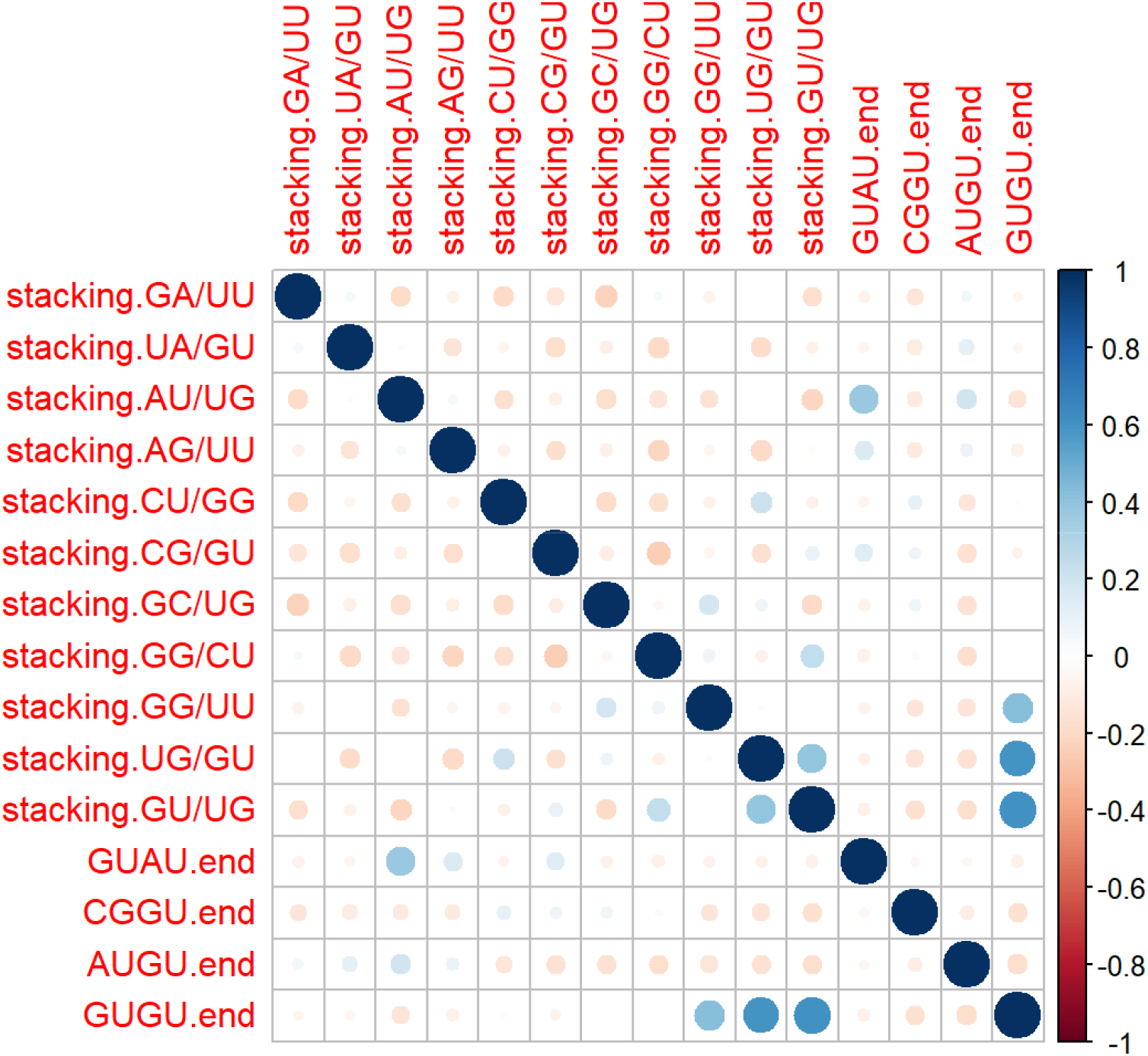
Pearson correlation between features of GU stacking model. Pairwise Pearson correlation coefficients were calculated for the observed frequencies of GU stacking parameters. The correlation coefficients are encoded in the size and color of each circle.

**Supplemental Figure 7:**
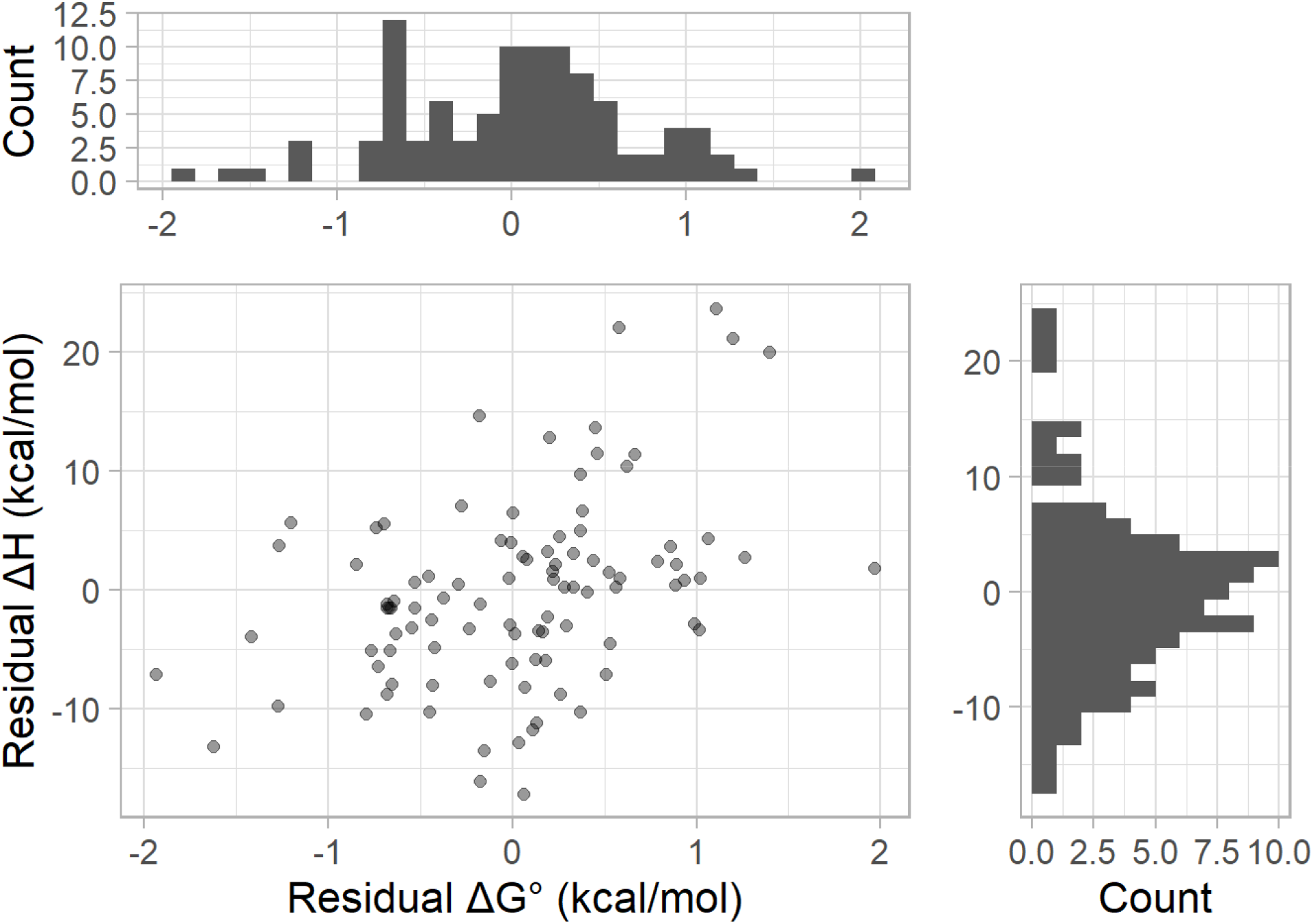
Residuals for the GU stacking parameter predictions. Shown are the scatter plot between the residuals in ΔH° and folding 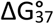 predictions (bottom left plot), a histogram of the folding 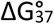 residuals (top plot), and a histogram of the ΔH° residuals right plot).

**Supplemental Figure 8:**
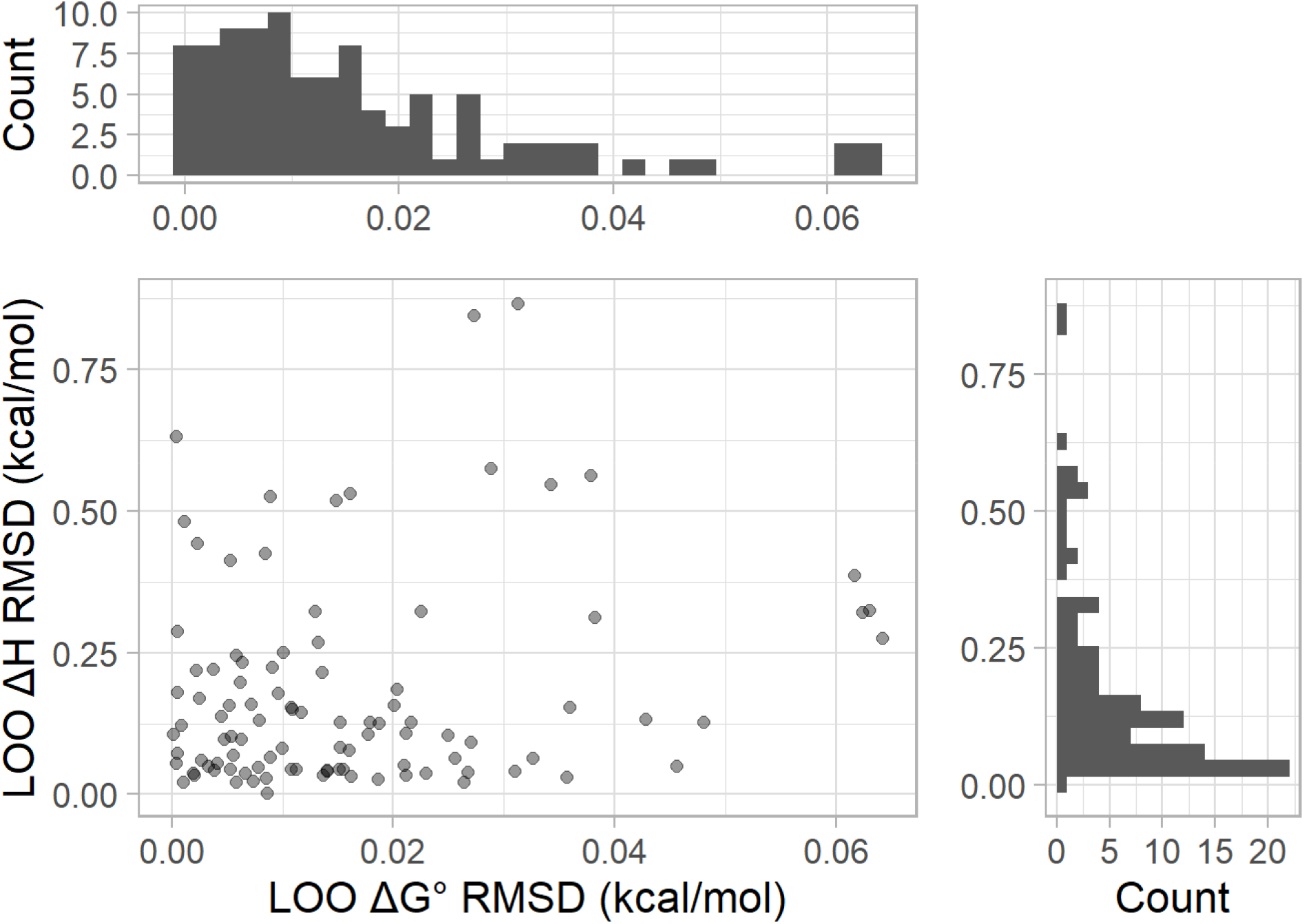
Leave-One-Out (LOO) analysis for GU stacking parameters. The plotted value is the RMSD of the parameter values for leaving out a single experiment as compared to when all experiments are used.

**Supplemental Figure 9.**
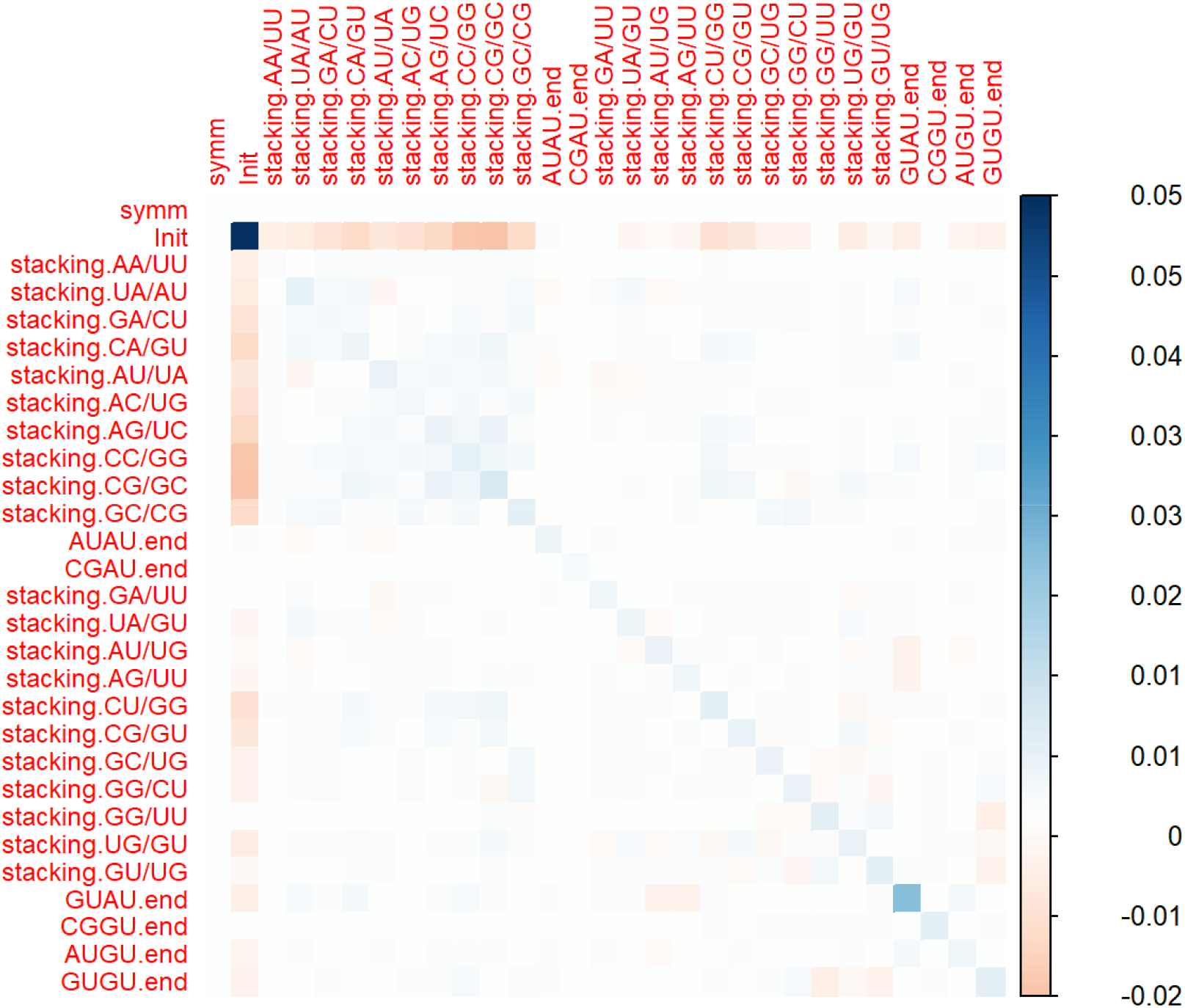
The covariation between 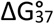 parameter values. Pairwise covariation values for 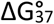 values fit to 1,000 randomly perturbed sets of optical melting experiments.

**Supplemental Figure 10.**
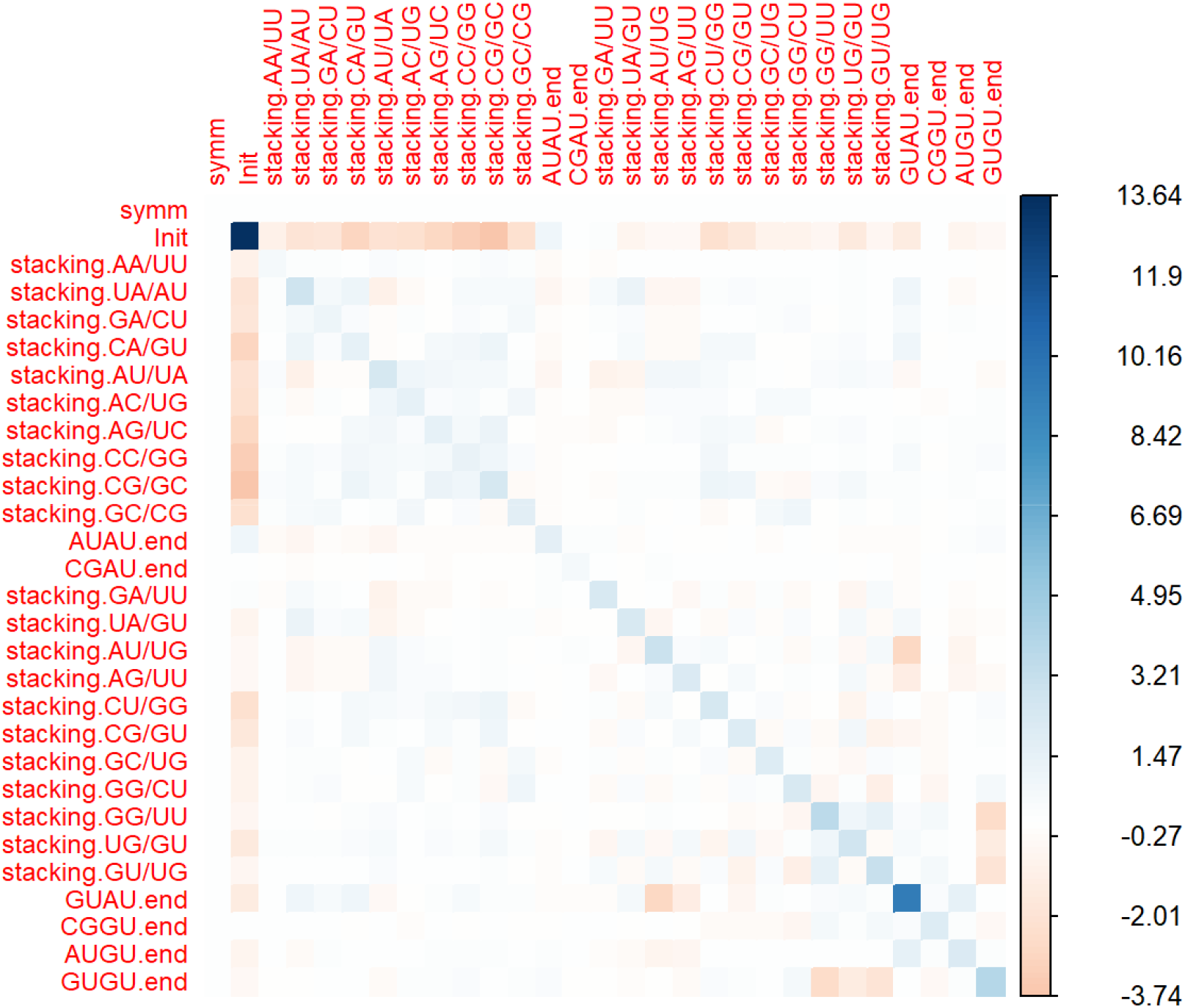
The covariation between ΔH° parameter values. Pairwise covariation values for ΔH° values fit to 1,000 randomly perturbed sets of optical melting experiments.

**Supplemental Figure 11:**
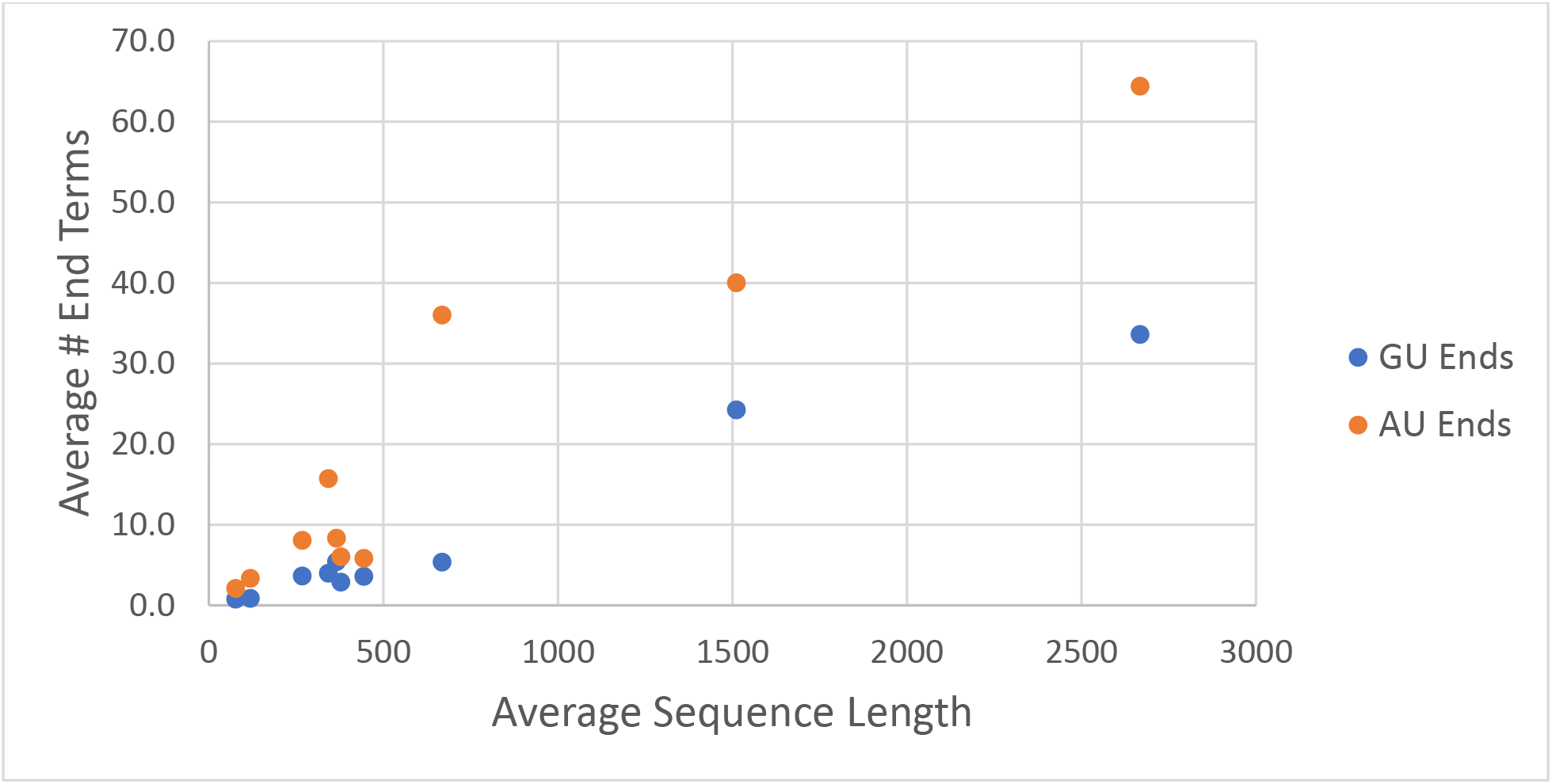
AU and GU helix end counts across RNA families. The accepted secondary structures of RNA sequences in different RNA families were parsed to count the number of AU and GU helix ends that existed in the structures. The RNA families included 5S rRNA, 16S rRNA, 23S rRNA, Group 1 and Group 2 introns, RNAP, SRP, telomerase, tmRNA, and tRNA. AU and GU ends were counted if they closed exterior loops, interior loops, hairpin loops, multibranch loops, or bulge loops larger than a single nucleotide.

